# Impact of a native hemiparasitic plant on invasive and native hosts in the field

**DOI:** 10.1101/2022.01.22.477372

**Authors:** Robert M. Cirocco, Jennifer R. Watling, José M. Facelli

## Abstract

There is increasing evidence that native parasitic plants are showing potential as biocontrols for plant invasions which are a major global threat to biodiversity. However, for this potential to be realised, the range of invasive hosts that can be controlled needs to be identified while also evaluating any potential off-target effects the parasite may pose. To address this, we investigated the impact of the Australian native shoot hemiparasitic plant, *Cassytha pubescens* on the major invasive species *Rubus anglocandicans* and two native shrubs, *Acacia pycnantha* and *Bursaria spinosa* in naturally occurring populations in southern Australia. We measured foliar predawn and midday quantum yield, electron transport rate (ETR), stomatal conductance (g_s_), carbon isotope composition and foliar nitrogen concentration [N] of uninfected and infected plants and, apart from g_s_, also for *C. pubescens*. Infection significantly decreased predawn and midday quantum yield, ETR and g_s_ of invasive *R. anglocandicans*. In contrast, infection had no effect on predawn and midday quantum yield, ETR or carbon isotope composition of the native hosts. However, *C. pubescens* had a significant negative effect on native host g_s_, and a positive effect on host [N]. Parasite stem [N] was significantly higher when infecting *A. pycnantha* than *B. spinosa*. These results strengthen evidence for native parasitic plants having greater impact on invasive hosts while having mild off-target effects on native hosts and thus, show potential to mitigate plant invasions and help protect biodiversity.

## Introduction

Parasitic plants can have profound effects on their hosts and also at population, community and landscape levels (Press and Phoenix 2005). For example, Hartley et al. (2015) found that the annual root hemiparasite *Rhinanthus minor* had a differential impact on grasses relative to forbs (tolerant host), resulting in a significant increase in plant diversity as well as invertebrate abundance. This may be due to parasite haustoria more effectively connecting to the vasculature of grasses than forbs (Cameron and Seel 2007). Similar differential effects on hosts have been observed where native parasitic plants infect introduced and native hosts in some ecosystems. This has led to suggestions that native parasites may have potential as novel biocontrols for weeds to help restore and conserve biodiversity (Těšitel et al. 2020). For instance, in China, the native parasitic vine *Cuscuta australis* was found to negatively impact growth of three major invasive plants resulting in a significant increase in native species cover and diversity (Yu et al. 2011). Li et al. (2015) also found that *Cuscuta australis* had a significant negative effect on total biomass of juvenile, invasive *Bidens pilosa*. Also in China, Li et al. (2012) found that the native *Cuscuta chinensis* caused significant damage to three invasive but not three native hosts.

In Australia, the native parasitic vine *Cassytha pubescens* has been shown to have a strong negative impact on the physiology and growth of the major invasive shrubs, *Ulex europaeus* and *Cytisus scoparius* (Prider et al. 2009; Shen et al. 2010; Cirocco et al. 2016a, b, 2017, 2018, 2020, 2021a). Moreover, *C. pubescens* has little effect on physiology and growth of the native hosts *Leptospermum myrsinoides* and *Acacia paradoxa* (Prider et al. 2009; Cirocco et al. 2016a, 2017; but see Cirocco et al. 2021b). This differential impact may be explained by *Cassytha’s* haustoria connecting more effectively to invasive than native hosts, facilitating greater resource removal from the former (Facelli et al. 2020).

For the true potential of *C. pubescens* to be realised, firstly, we must understand if its greater effects on invasive hosts extend to other species, particularly those which are most problematic. *Cassytha pubescens* has been observed infecting one of Australia’s worst weeds, European blackberry (*Rubus fruticosus* agg.), which is also a major invasive species in parts of North America, Hawaii, Chile, and was declared the worst weed in New Zealand (Parsons and Cuthbertson 2001). European blackberry has very few classical biocontrol options available apart from a leaf rust fungus, *Phragmidium violaceum* that has had variable success (Amor et al. 1998). We also need to determine if *C. pubescens* has a negative impact on native hosts not yet studied, to identify ‘off-target’ impacts that may make it unsuitable for biocontrol in certain situations. This is important, particularly as recent evidence suggests that in some instances, native hosts may be adversely affected by the parasite (Cirocco et al. 2021b). There is also an urgent need for identifying effective weed biocontrols considering that plant invasions are a major threat to global diversity (Vilà et al. 2019).

We are also interested in gaining a mechanistic understanding of the differential impacts of native parasites on native and introduced hosts. It is possible that native parasites more effectively remove resources from invasive hosts, leading to them becoming water and nutrient stressed. Stressed hosts with decreased stomatal conductance would also have lower rates of photosynthesis, which at a constant or increasing photosynthetic photon flux density (PPFD) would expose infected plants to conditions of excess light (Demmig-Adams and Adams 1992). Prolonged exposure to excess light could lead to increased susceptibility to chronic photoinhibition, reflected by declines in *F*_v_/*F*_m_ (Demmig-Adams and Adams 2006; Cirocco et al. 2016b, 2021a). Prolonged impacts on these higher order physiological traits will ultimately lead to lower growth, reproductive output, and the possible death of the host. As *C. pubescens* is a perennial, vine with indeterminate growth, it can, and does, attach to multiple hosts, and when nearby hosts are reachable and susceptible to infection, the parasite is not impacted by the death of an individual host.

Here we report results of field studies on the impact of *C. pubescens* on invasive *R. anglocandicans* and two native hosts, *Acacia pycnantha* and *Bursaria spinosa*. We hypothesised that the native parasite would have a negative impact on a range of physiological traits in the invasive host, but not the two native hosts. To determine parasite impact, we measured stomatal conductance (g_s_), stable carbon isotope composition (δ^13^C), foliar nitrogen concentration [N], photosynthesis and photoinhbition of both uninfected and infected plants. We also measured the performance of *C. pubescens*, using these same variables (apart from g_s_).

## Materials and methods

### Study species

*Cassytha pubescens* (Lauraceae) is a native Australian, perennial, hemiparasitic vine that has stems 0.5–1.5 mm in diameter but no true leaves (McLuckie 1924; Harden 1990; Kokubugata et al. 2012). It coils around host shoots, forming multiple haustoria 2–3 mm in diameter (McLuckie 1924; Harden 1990). It occurs throughout south-eastern Australia and is a generalist parasite often infecting both native, and invasive, perennial shrubs (McLuckie 1924; Harden 1990; Cirocco et al. 2018).

*Rubus anglocandicans* (Rosaceae) which is the most prevalent member of the *Rubus fruticosus agg*. in Australia (Evans and Weber 2003), is a perennial shrub that commonly forms impenetrable thickets 2–3 m high (Amor et al. 1998; Evans et al. 2007). In Australia, *R. anglocandicans* grows in a variety of soil types and where annual rainfall is higher than 760 mm or water is readily available (e.g. creeks) (Amor et al. 1998). It is a highly invasive species due to its vigorous growth, ability to reproduce both vegetatively and sexually, and effective, long distance seed dispersal via fruitivores or water systems (Amor et al. 1998; Evans et al. 2007). *Acacia pycnantha* (Fabaceae) is a perennial, evergreen shrub or small tree that can grow to approx. 5 m and flowers in spring (Spencer 2002). It is native to south-eastern Australia but has become invasive in Western Australia, South Africa and Portugal (Ndlovu et al. 2013). *Bursaria spinosa* (Pittosporaceae) is a perennial, evergreen shrub or small tree that is 1–3 m tall but can reach approx. 9 m (Cayzer et al. 1999; Cunningham et al. 2011). Shoots are spiny particularly when young and flowering occurs mainly in summer (Cayzer et al. 1999).

### Description of studies

From the outset we emphasise the field studies described below were conducted in different years and across different locations and thus it was appropriate to analyse them separately. They were, however, conducted in the same region and vegetation types on uninfected and infected hosts of similar size (1–3 m high) with infected plants having a similar level of parasite present (about 50% of the host covered with *C. pubescens*) (Suppl. Figs S1–S2).

All studies were conducted in the Mt Lofty Ranges of South Australia which cover an area of 6,282 km^2^. Only 13% of the native vegetation remains in this region, which comprises a eucalypt woodland overstorey, an understorey of mainly sclerophyllous shrubs and a ground layer of grasses and sedges (Armstrong et al. 2003). The climate is Mediterranean, with hot dry summers and cold wet winters (Armstrong et al. 2003). The invasive host, *R. anglocandidans* was studied at two sites, Horsnell Gully (34°93’53”S, 138°70’45”E) and Morialta (34°90’55”S, 138°71’22”E) Conservation Parks, where it occurs, either uninfected or naturally infected with *C. pubescens*. Ecophysiological measurements were made on 9–12 uninfected and infected canes of *R. anglocandicans* at each site and also on the parasite. Individual canes were considered single replicates based on McDowell (2002), and unpublished work in our lab that found parasite effects were localised to infected canes of this species.

The native hosts were studied at two sites within Belair National Park: Queens Jubilee Drive (35°0’33”S, 138°40’14”E) and Saddle Hill (35°0’40”S, 138°40’37”E), where they occur either uninfected or naturally infected with *C. pubescens*. Ecophysiological measurements were made on 6–7 uninfected and infected plants at each site. All measurements below were made on the youngest fully expanded leaf or phyllode (in the case of *A. pycnantha*).

### Host and parasite chlorophyll fluorescence and host stomatal conductance

Predawn (*F*_v_/*F*_m_) and midday (Φ_PSII_) quantum yields and midday electron transport rates (ETR) were measured with a MINI-PAM fitted with a 2030–B leaf-clip (Walz, Effeltrich, Germany). *R. anglocandidans* was measured in mid-April 2019, and *A. pycnantha* and *B. spinosa* in late-February 2020. Measurements were conducted on sunny days between 12:00-15:00. Mean PPFD (μmol m^−2^s^−1^) was 1221 ± 3 for the invasive host and parasite across the two sites (*n* = 66), and 1481 ± 4 for native host species and parasite across the two sites (*n* = 123) (Suppl. Figs S3–S4).

Stomatal conductance (g_s_) was measured with a portable, SC-1 Leaf Porometer (Decagon Devices, Inc. Washington) in late-April 2019 for *R. anglocandidans* and early March 2020 for the two native hosts. Measurements were made between 09:00-12:00 on sunny days (Suppl. Figs S3–S4). As, the porometer only measures flat leaves, g_s_ of parasite stems could not be measured.

### Host and parasite δ^13^C and nitrogen concentration

Host foliage and parasite stems were collected at the same time as physiological measurements, and were dried at 60 °C for seven days and then finely ground. Carbon isotope composition was determined by mass spectrometry and nitrogen concentration [N] by elemental analyser using either an IsoPrime isotope ratio mass spectrometer (GV Instruments, Manchester, UK) and Elementar Isotope CUBE Elemental Analyser (Elementar Analysensysteme, Hanau, Germany) for the parasite:invasive host association, or a Nu Horizon isotope ratio mass spectrometer, (Wrexham, UK) equipped with an elemental analyser (EA3000, EuroVector, Pavia, Italy) for the parasite:native host associations.

### Statistical analyses

The effect of infection and site on physiological performance of *R. anglocandicans* was analysed with a two-way, fixed effects ANOVA (sites not chosen at random). The effect of site on parasite performance was analysed with a one-way, fixed effects ANOVA. For the native hosts, the effect of infection, host species and site on uninfected and infected plants (infected shoots) was analysed using a three-way ANOVA (fixed effects: sites not chosen at random). The effects of host species and site on parasite data were analysed with a two-way ANOVA. Any significant interactions were subjected to pairwise comparisons within each level of the other effect(s) in the model using a Tukey HSD test. When an interaction was not detected, main effects became valid. Model assumptions were met, in some cases this required data transformation. All data were analysed in R (R Development Core Team, 2016) and *α* = 0.05 (Type 1 error rate), and graphs were generated with GraphPad prism (La Jolla, CA, USA).

## Results

### R. anglocandidans

Infection had a main effect on *F*_v_/*F*_m_, Φ_PSII_ and ETR of *R. anglocandicans* (Table 1; no interactions: Fig. 1a, d, f). *F*_v_/*F*_m_, Φ_PSII_ and ETR of infected plants were 3%, 23% and 22% lower than uninfected plants, respectively (Fig. 1b, e, g). Host *F*_v_/*F*_m_ also differed significantly between sites and was 2% lower at Horsnell than that at Morialta (Table 1; Fig. 1c). Site had no effect on parasite *F*_v_/*F*_m_, Φ_PSII_ or ETR (Table 1; Fig. S5a, b, c). Main effects of infection and site were found for host g_s_ (Table 1; no interaction: Fig. 2a). Stomatal conductance of infected plants was lower by 21% relative to that of uninfected plants (Fig. 2b). Host g_s_ at Horsnell was 34% lower than that at Morialta (Fig. 2c).

**Table 1.** ANOVA results for 1) the effects of *Cassytha pubescens* infection (I) and site (S) on foliar predawn and midday quantum yield (*F*_v_/*F*_m_ and Φ_PSII_), midday electron transport rates (ETR), stomatal conductance (g_s_), nitrogen concentration [N] and carbon isotope composition (δ^13^C) of *Rubus anglocandicans* at two field sites and 2) Site effects on *F*_v_/*F*_m_, Φ_PSII_, ETR and δ^13^C of *C. pubescens* when infecting *R. anglocandicans* at these sites

| (Host) | $F_v/F_m$ | $\Phi_{PSII}$ | ETR | $g_s$ | [N] | $\delta^{13}C$ |
| --- | --- | --- | --- | --- | --- | --- |
| I | **0.0003\*\*\*** | **0.005\*\*** | **0.007\*\*** | **0.052\*** | **0.110** | **0.073** |
|  | *15.4* | *8.78* | *8.08* | *4.04* | *2.68* | *3.42* |
|  | 0.008 | 0.019 | 4701 | 21.5 | 0.210 | 1.53 |
| S | **0.009\*\*** | **0.495** | **0.442** | **0.001\*\*** | **0.561** | **0.024\*** |
|  | *7.58* | *0.475* | *0.603* | *12.4* | *0.345* | *5.55* |
|  | 0.004 | 0.001 | 351 | 65.9 | 0.027 | 2.48 |
| I × S | **0.930** | **0.277** | **0.304** | **0.378** | **0.361** | **0.252** |
|  | *0.008* | *1.21* | *1.09* | *0.796* | *0.857* | *1.35* |
|  | 0.000004 | 0.003 | 632 | 4.24 | 0.067 | 0.605 |
| Error | 0.023 | 0.087 | 23281 | 192 | 2.82 | 16.1 |
| *df* | 1, 44 | 1, 40 | 1, 40 | 1, 36 | 1, 36 | 1, 36 |
| (Parasite) | $F_v/F_m$ | $\Phi_{PSII}$ | ETR | $g_s$ | [N] | $\delta^{13}C$ |
| S | **0.970** | **0.175** | **0.199** | N/A | N/A | **<0.0001\*\*\*** |
|  | *0.002* | *1.98* | *1.76* |  |  | *29.8* |
|  | 0.0005 | 0.017 | 3981 |  |  | 18.3 |
| Error | 8.15 | 0.175 | 45184 | N/A | N/A | 11.0 |
| *df* | 1, 22 | 1, 20 | 1, 20 | N/A | N/A | 1, 18 |

*P*, *F* and sum of square values are in **bold**, *italic* and regular type, respectively. Degrees of freedom = *df*, and not applicable = N/A. Significance codes: <0.0001-0.001'\*\*\*', 0.001-0.01'\*\*, 0.01-0.05'\*. Host $g_s$ and parasite $F_v/F_m$ square root and inverse log transformed, respectively. Welch two sample *t*-test used for parasite [N] as data did not meet assumptions of ANOVA: $t_{11,2} = 0.608$

**Fig. 1.**
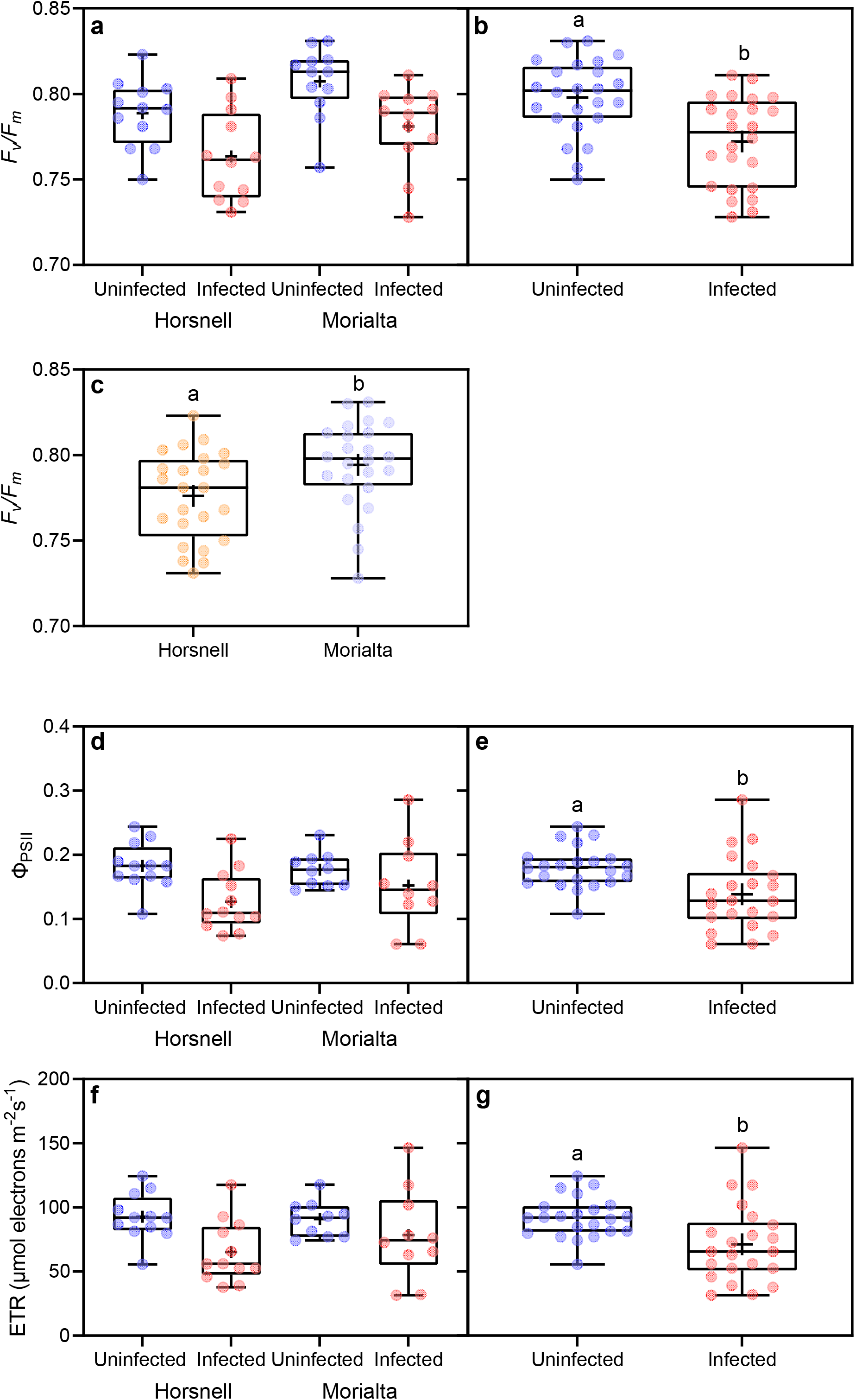
(**a**) Predawn (*F*_v_/*F*_m_) for *Rubus anglocandicans* either uninfected or infected with *Cassytha pubescens* at two sites (Horsnell or Morialta) in the Mt. Lofty Ranges of South Australia, and main infection (**b**) and site (**c**) effects. Similarly, midday (Φ_PSII_) quantum yield (**d**), and main infection effect (**e**), and midday electron transport rates (ETR) (**f**), and main infection effect (**g**). All data points, median, 1^st^ and 3^rd^ quartiles, interquartile range and mean (+ within box) are shown, different letters signify significant differences: *n* = 12 (**a**), *n* = 10–12 (**d**, **f**), *n* = 24 (**b**, **c**) and *n* = 22 (**e**, **g**)

**Fig. 2.**
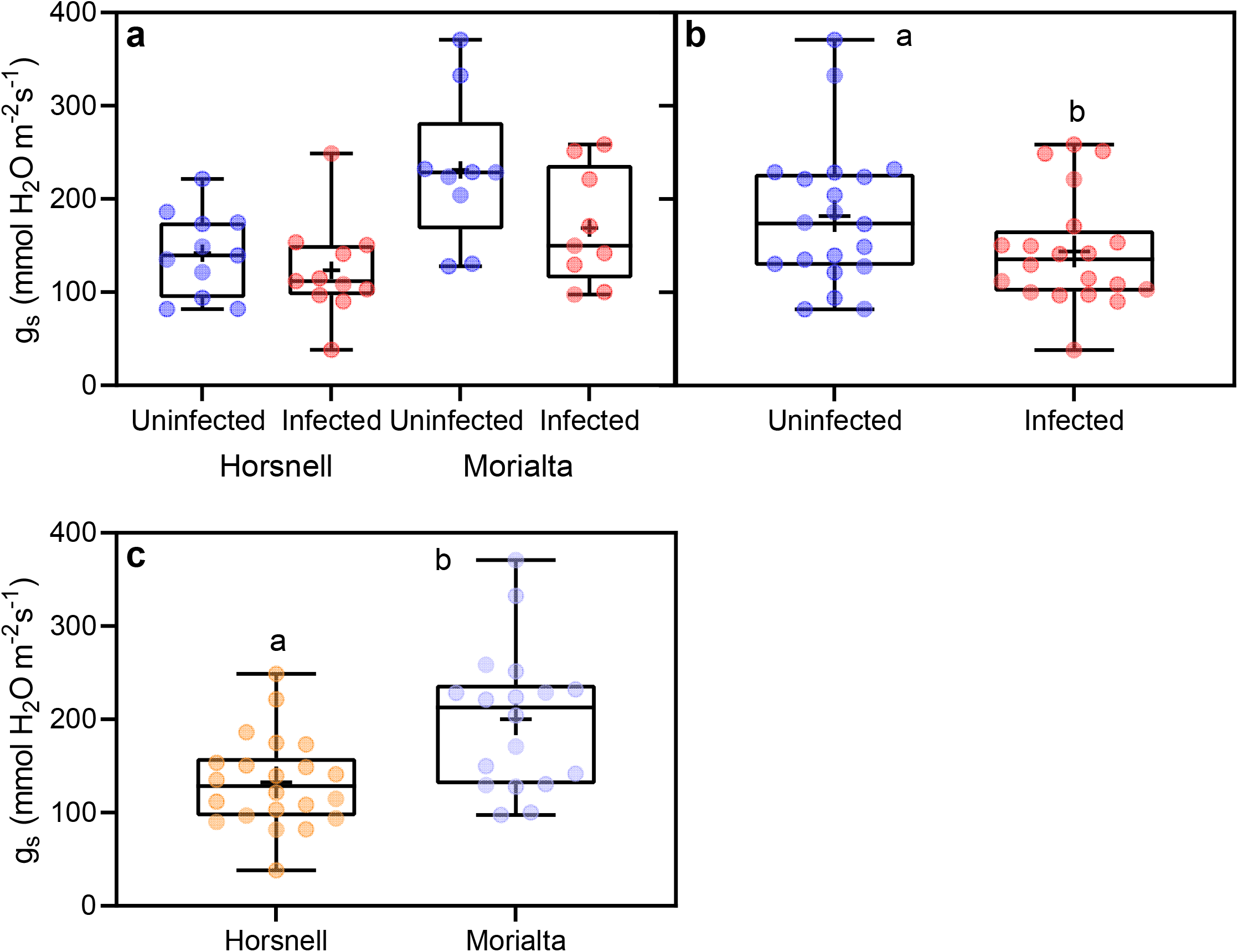
(**a**) Stomatal conductance (g_s_) of *Rubus anglocandicans* either uninfected or infected with *Cassytha pubescens* at two sites (Horsnell or Morialta). Main effect of (**b**) infection and (**c**) site on host g_s_. All data points, median, 1^st^ and 3^rd^ quartiles, interquartile range and mean (+ within box) are shown, different letters signify significant differences and *n* = 9–11 (**a**), *n* = 20 (**b**, **c**)

No significant infection or site effects were found for leaf [N] of *R. anglocandicans* (Table 1; Fig. 3a). Site had no effect on parasite stem [N] (*P* = 0.555) (Table 1; Fig. 3b). There was a main site effect on δ^13^C of *R. anglocandicans* (Table 1; no interaction: Fig. 4a). Host δ^13^C at Horsnell was significantly higher than that at Morialta (Fig. 4b). Site also significantly affected parasite stem δ^13^C, which was significantly higher at Horsnell than at Morialta (Table 1; Fig. 4c). Comparing δ^13^C of infected *R. anglocandicans* and associated *C. pubescens* across the two sites, a significant species × site interaction was detected (*F*_1, 36_ = 14.6; *P* = 0.0005). δ^13^C of *C. pubescens* was significantly higher than that of *R. anglocandicans* at Horsnell while there was no difference detected at Morialta (Fig. 4c).

**Fig. 3.**
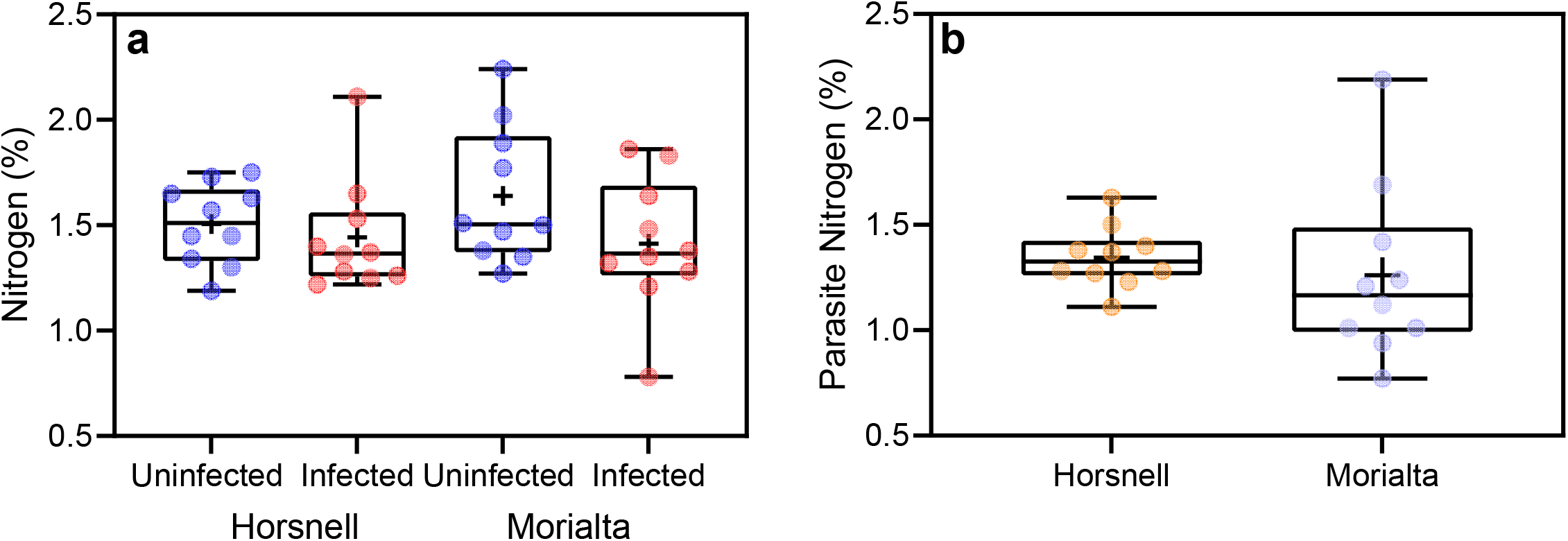
(**a**) Foliar nitrogen concentration [N] of *Rubus anglocandicans* either uninfected or infected with *Cassytha pubescens* at two sites (Horsnell or Morialta). (b) Stem [N] concentration of *C. pubescens* when infecting *R. anglocandicans* at Horsnell or Morialta. All data points, median, 1^st^ and 3^rd^ quartiles, interquartile range and mean (+ within box) are shown, different letters signify significant differences and *n* = 10 (**a**, **b**)

**Fig. 4.**
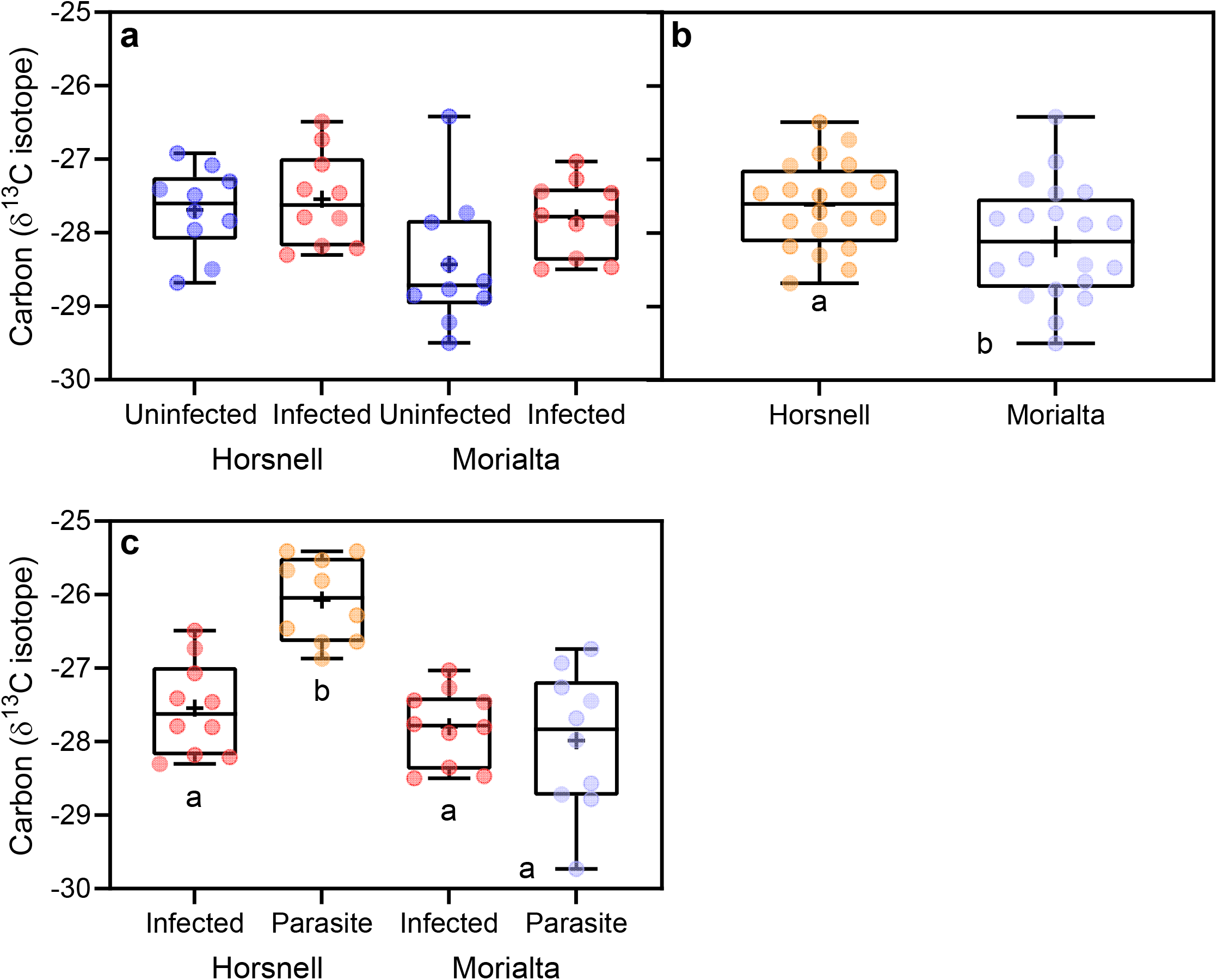
(**a**) Foliar carbon isotope composition (δ^13^C) of *Rubus anglocandicans* either uninfected or infected with *Cassytha pubescens* at two sites (Horsnell or Morialta). (**b**) Main effect of site on host δ^13^C. (**c**) δ^13^C of infected *R. anglocandicans* (Infected) and *C. pubescens* (Parasite), at Horsnell (H) or Morialta (M). All data points, median, 1^st^ and 3^rd^ quartiles, interquartile range and mean (+ within box) are shown, different letters signify significant differences: *n* = 10 (**a**, **c**), *n* = 20 (**b**)

### Native hosts

A species × site interaction was found for host *F*_v_/*F*_m_ (Table 2; no three-way interaction: Fig 5a). *F*_v_/*F*_m_ of *A. pycnantha* was significantly higher at Queens Jubilee than Saddle Hill, while *F*_v_/*F*_m_ of *B. spinosa* did not differ between sites (Fig. 5b). Site had a main effect on host Φ_PSII_ and ETR (Table 2; no interactions: Fig. 5c, e). Host Φ_PSII_ and ETR at Queens Jubilee were approx. 25% higher compared with those at Saddle Hill (Fig. 5d, f). *F*_v_/*F*_m_ of *C. pubescens* (when infecting *A. pycnantha*), was significantly higher at Queens Jubilee than Saddle Hill, and also higher than that of *C. pubescens* infecting *B. spinosa* at either site (Table 3; Fig. S6a). Site had a main effect on parasite Φ_PSII_ and ETR (Table 3; no interactions: Fig S6b, d). Parasite Φ_PSII_ and ETR at Queens Jubilee were approx. 30% higher than at Saddle Hill (Fig S6c, e). A species × site and main effect of infection were found for host g_s_, (Table 2; no three-way interaction: Fig. 6a). Stomatal conductance of *B. spinosa* at Saddle Hill was significantly higher than at Queens Jubilee, and also than that of *A. pycnantha* at both sites (Fig. 6b). Stomatal conductance of infected plants was 17% lower than that of uninfected ones (Fig. 6c).

**Table 2.**
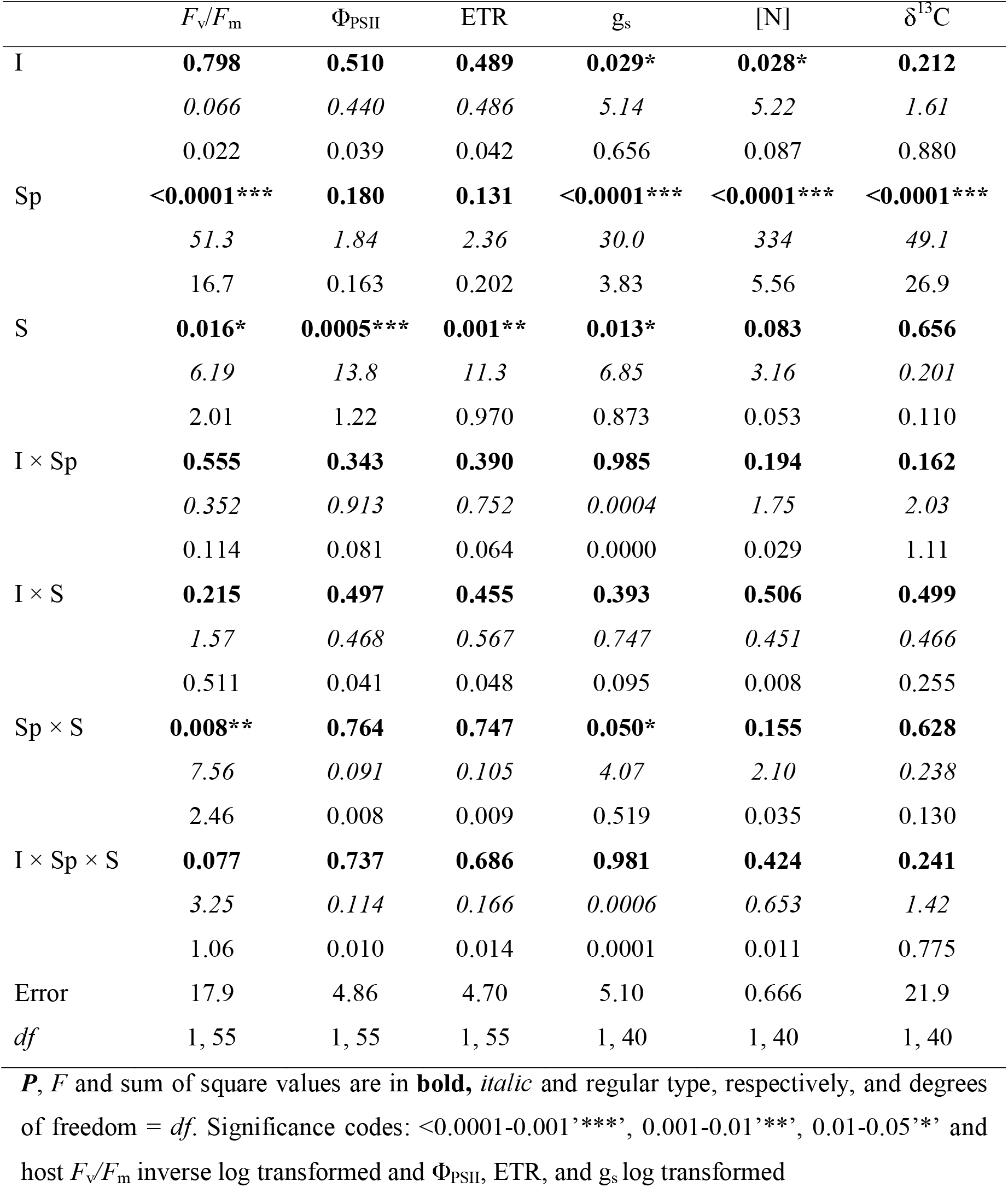
ANOVA results for the effects of *Cassytha pubescens* infection (I), host species (Sp) and site (S) on foliar predawn and midday quantum yield (*F*_v_/*F*_m_ and Φ_PSII_), midday electron transport rates (ETR), stomatal conductance (g_s_), nitrogen concentration [N] and carbon isotope composition (δ^13^C) of *Acacia pycnantha* or *Bursaria spinosa* at two sites

| | $F_v/F_m$ | $\Phi_{PSII}$ | ETR | $g_s$ | [N] | $\delta^{13}C$ |
| --- | --- | --- | --- | --- | --- | --- |
| I | **0.798** | **0.510** | **0.489** | **0.029\*** | **0.028\*** | **0.212** |
|  | *0.066* | *0.440* | *0.486* | *5.14* | *5.22* | *1.61* |
|  | 0.022 | 0.039 | 0.042 | 0.656 | 0.087 | 0.880 |
| Sp | **<0.0001\*\*\*** | **0.180** | **0.131** | **<0.0001\*\*\*** | **<0.0001\*\*\*** | **<0.0001\*\*\*** |
|  | *51.3* | *1.84* | *2.36* | *30.0* | *334* | *49.1* |
|  | 16.7 | 0.163 | 0.202 | 3.83 | 5.56 | 26.9 |
| S | **0.016\*** | **0.0005\*\*\*** | **0.001\*\*** | **0.013\*** | **0.083** | **0.656** |
|  | *6.19* | *13.8* | *11.3* | *6.85* | *3.16* | *0.201* |
|  | 2.01 | 1.22 | 0.970 | 0.873 | 0.053 | 0.110 |
| I × Sp | **0.555** | **0.343** | **0.390** | **0.985** | **0.194** | **0.162** |
|  | *0.352* | *0.913* | *0.752* | *0.0004* | *1.75* | *2.03* |
|  | 0.114 | 0.081 | 0.064 | 0.0000 | 0.029 | 1.11 |
| I × S | **0.215** | **0.497** | **0.455** | **0.393** | **0.506** | **0.499** |
|  | *1.57* | *0.468* | *0.567* | *0.747* | *0.451* | *0.466* |
|  | 0.511 | 0.041 | 0.048 | 0.095 | 0.008 | 0.255 |
| Sp × S | **0.008\*\*** | **0.764** | **0.747** | **0.050\*** | **0.155** | **0.628** |
|  | *7.56* | *0.091* | *0.105* | *4.07* | *2.10* | *0.238* |
|  | 2.46 | 0.008 | 0.009 | 0.519 | 0.035 | 0.130 |
| I × Sp × S | **0.077** | **0.737** | **0.686** | **0.981** | **0.424** | **0.241** |
|  | *3.25* | *0.114* | *0.166* | *0.0006* | *0.653* | *1.42* |
|  | 1.06 | 0.010 | 0.014 | 0.0001 | 0.011 | 0.775 |
| Error | 17.9 | 4.86 | 4.70 | 5.10 | 0.666 | 21.9 |
| df | 1, 55 | 1, 55 | 1, 55 | 1, 40 | 1, 40 | 1, 40 |

**P**, **F** and sum of square values are in **bold**, *italic* and regular type, respectively, and degrees of freedom = *df*. Significance codes: <0.0001-0.001'\*\*\*', 0.001-0.01'\*\*\*', 0.01-0.05'\*' and host $F_v/F_m$ inverse log transformed and $\Phi_{PSII}$ , ETR, and $g_s$ log transformed

**Fig. 5.**
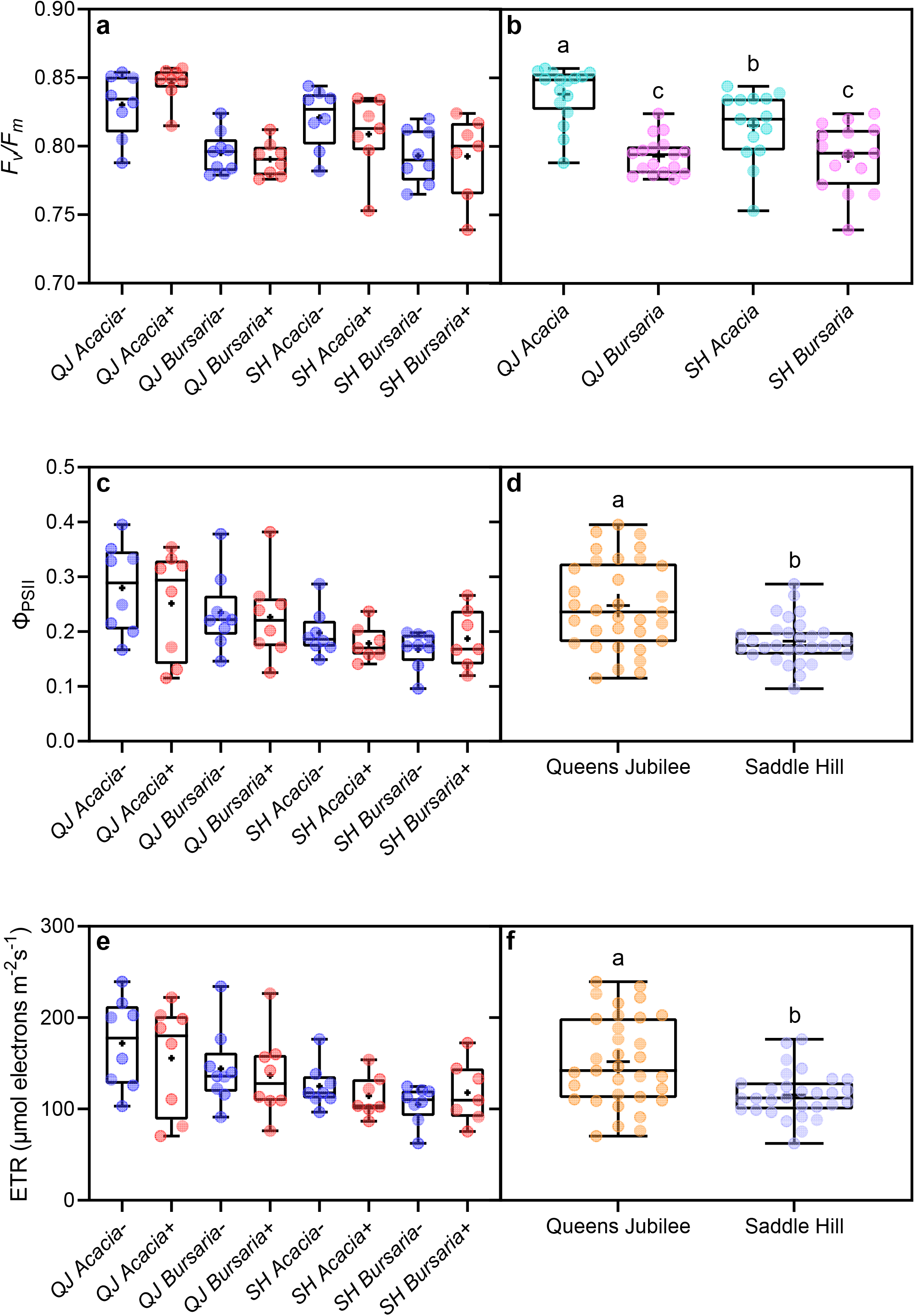
(**a**) Predawn (*F*_v_/*F*_m_) quantum yield for *Acacia pycnantha* and *Bursaria spinosa* either uninfected (−) or infected (+) with *Cassytha pubescens* at two sites: Queens Jubilee (QJ) or Saddle Hill (SH), and (**b**) Species × site effect on host *F*_v_/*F*_m_. Similarly, (**c**) midday (Φ_PSII_) quantum yield, and (**d**) main effect of site on host Φ_PSII_, and (**e**) midday electron transport rates, and (**f**) main effect of site on host Φ_PSII_. All data points, median, 1^st^ and 3^rd^ quartiles, interquartile range and mean (+ within box) are shown, different letters signify significant differences: *n* = 7–9 (**a**, **c**, **e**), *n* = 15–17 (**b**) and *n* = 30–33 (**d**, **f**)

**Table 3.** ANOVA results for the effects of species (Sp) and site (S) on stem predawn and midday quantum yield (*F*_v_/*F*_m_ and Φ_PSII_), midday electron transport rates (ETR), stomatal conductance (g_s_), nitrogen concentration [N] and carbon isotope composition (δ^13^C) of *C. pubescens* when infecting *Acacia pycnantha* or *Bursaria spinosa* at two field sites (Queens Jubilee and Saddle Hill). Also includes comparison between δ^13^C of parasite and *A. pycnantha* or *B. spinosa* at the two sites

| | $F_v/F_m$ | $\Phi_{PSII}$ | ETR | [N] | $\delta^{13}C$ | $\delta^{13}C$<br>A.<br>*pycnantha* | $\delta^{13}C$<br>B.<br>*spinosa* |
| --- | --- | --- | --- | --- | --- | --- | --- |
| Sp | **0.181** | **0.312** | 0.278 | **0.0001\*\*\*** | **0.363** | **<0.0001\*\*\*** | **0.010\*\*** |
|  | *1.89* | *1.06* | *1.23* | *22.2* | *0.867* | *135* | *8.05* |
|  | 0.530 | 0.005 | 0.126 | 2.22 | 0.260 | 43.5 | 2.80 |
| S | **0.282** | **0.008\*\*** | **0.009\*\*** | **0.505** | **0.083** | **0.035\*** | **0.785** |
|  | *1.21* | *8.37* | *7.90* | *0.462* | *3.33* | *5.14* | *0.077* |
|  | 0.339 | 0.035 | 0.812 | 0.046 | 1.00 | 1.65 | 0.027 |
| Sp × S | **0.021\*** | **0.121** | *0.139* | **0.088** | **0.004\*\*** | **0.014\*** | **0.281** |
|  | *6.04* | *2.57* | *2.33* | *3.23* | *11.0* | *7.29* | *1.23* |
|  | 1.70 | 0.011 | 0.240 | 0.323 | 3.30 | 2.34 | 0.427 |
| Error | 7.30 | 0.110 | 2.67 | 2.00 | 6.01 | 6.43 | 6.96 |
| df | 1, 26 | 1, 26 | 1, 26 | 1, 20 | 1, 20 | 1, 20 | 1, 20 |

**P**, **F** and sum of square values are in **bold**, *italic* and regular type, respectively, and degrees of freedom = *df*. Significance codes: <0.0001-0.001'\*\*\*', 0.001-0.01'\*\*\*', 0.01-0.05'\*\*\*' and $F_v/F_m$ inverse log transformed and $\Phi_{PSII}$ , ETR, transformed to the power of 0.25

**Fig. 6.**
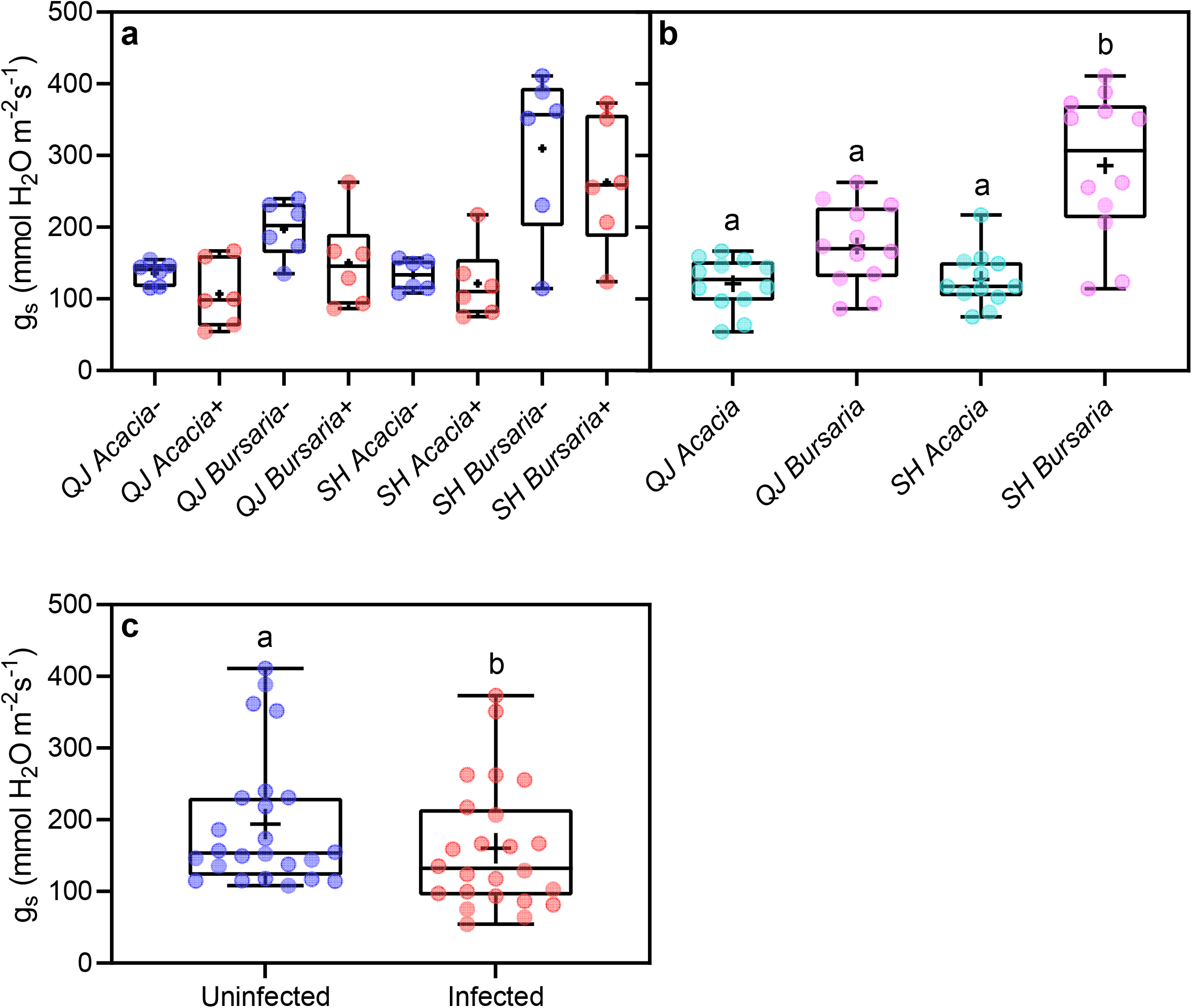
(**a**) Stomatal conductance (g_s_) of *Acacia pycnantha* and *Bursaria spinosa* either uninfected (–) or infected (+) with *Cassytha pubescens* at two sites: Queens Jubilee (QJ) or Saddle Hill (SH). (**b**) Species × site effect on g_s_ of *A. pycnantha* and *B. spinosa*. (**c**) Main effect of infection on host g_s_. All data points, median, 1^st^ and 3^rd^ quartiles, interquartile range and mean (+ within box) are shown, different letters signify significant differences and *n* = 6 (**a**), *n* = 12 (**b**) and *n* = 24 (**c**)

There were main effects of infection and species on host foliar [N] (Table 2; no interactions: Fig. 7a). Foliar [N] of infected plants was 6% higher than that of uninfected ones (Fig. 7b). Foliar N of *A. pycnantha* was 39% higher than *B. spinosa* (Fig. 7c). Species had a main effect on stem [N] of *C. pubescens* (Table 3; no interaction: Fig. 7d), which was 36% higher when infecting *A. pycnantha* than when infecting *B. spinosa* (Fig. 7e). Species had a main effect on host δ^13^C (Table 2; no interaction: Fig. 8a), which was significantly lower for *A. pycnantha* than *B. spinosa*, regardless of site or infection (Fig. 8b). δ^13^C of *C. pubescens* was significantly higher when growing on *A. pycnantha* than *B. spinosa* at Queens Jubilee, and also significantly higher when growing on *A. pycnantha* at Queens Jubilee than Saddle Hill (Table 3; Fig. 8c). Comparing δ^13^C of infected *A. pycnantha* and associated *C. pubescens* across the two sites, a significant species × site interaction was detected (Table 3), with δ^13^C of *C. pubescens* being significantly higher than that of *A. pycnantha* at both sites, but especially so at Queens Jubilee (Fig. 8d). Comparing δ^13^C of infected *B. spinosa* and associated *C. pubescens* across the two sites, a significant species effect was found (Table 3; no interaction: Fig. 8e), with δ^13^C of *C. pubescens* (–28.5 ‰ ± 0.2) being significantly higher than that of *B. spinosa* (–27.8 ‰ ± 0.2), regardless of site.

**Fig. 7.**
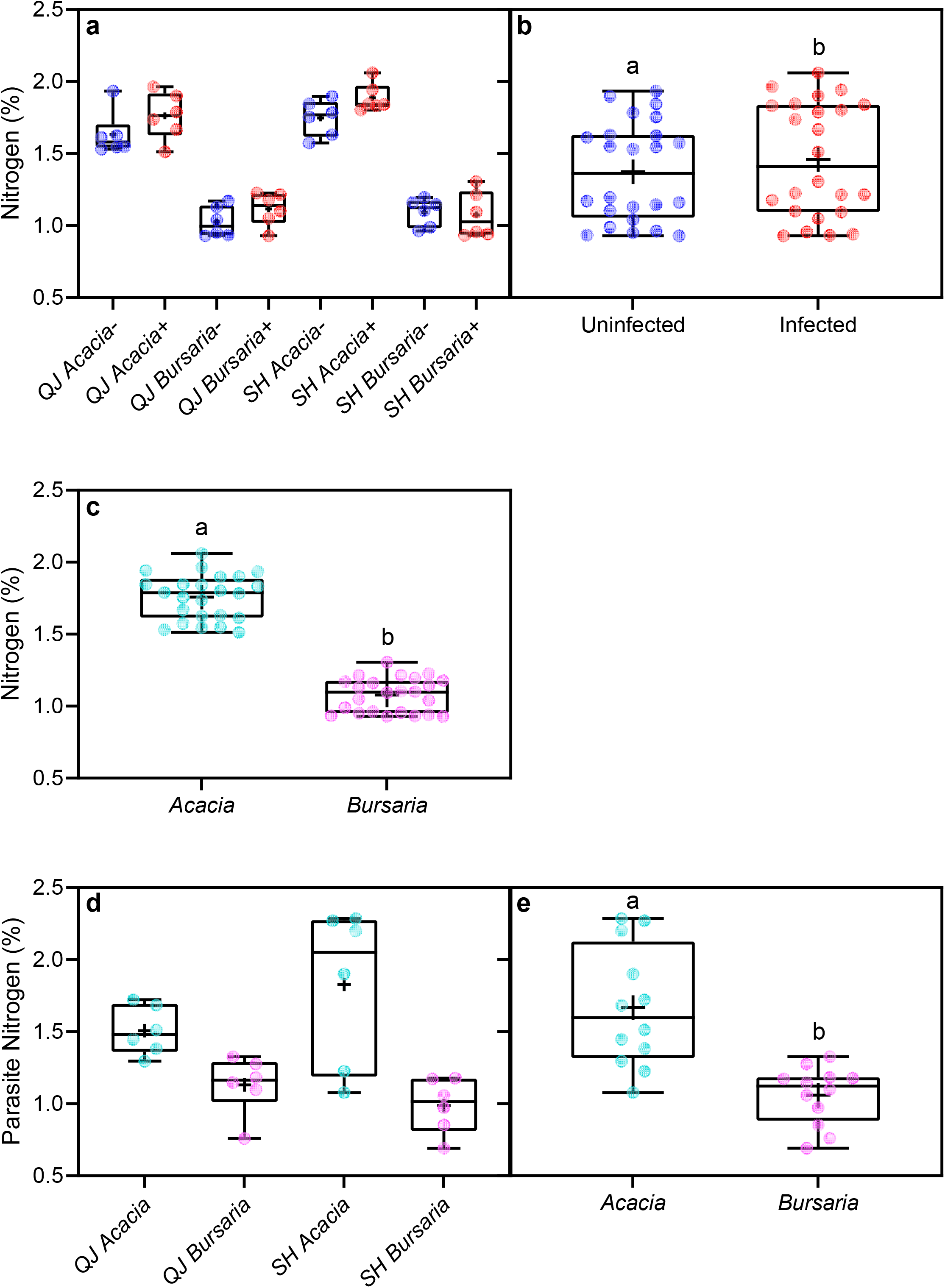
(**a**) Foliar nitrogen concentration [N] of *Acacia pycnantha* and *Bursaria spinosa* either uninfected (–) or infected (+) with *Cassytha pubescens* at two sites: Queens Jubilee (QJ) or Saddle Hill (SH). (**b**) Main effect of infection on host foliar N. (**c**) Main effect of species on host [N]. (**d**) Parasite [N] when infecting *Acacia pycnantha* or *Bursaria spinosa* at QJ or SH. (**e**) Main effect of species on parasite [N]. All data points, median, 1^st^ and 3^rd^ quartiles, interquartile range and mean (+ within box) are shown, different letters signify significant differences *n* = 6 (**a**, **d**), *n* = 24 (**b**, **c**) and *n* = 12 (**e**)

**Fig. 8.**
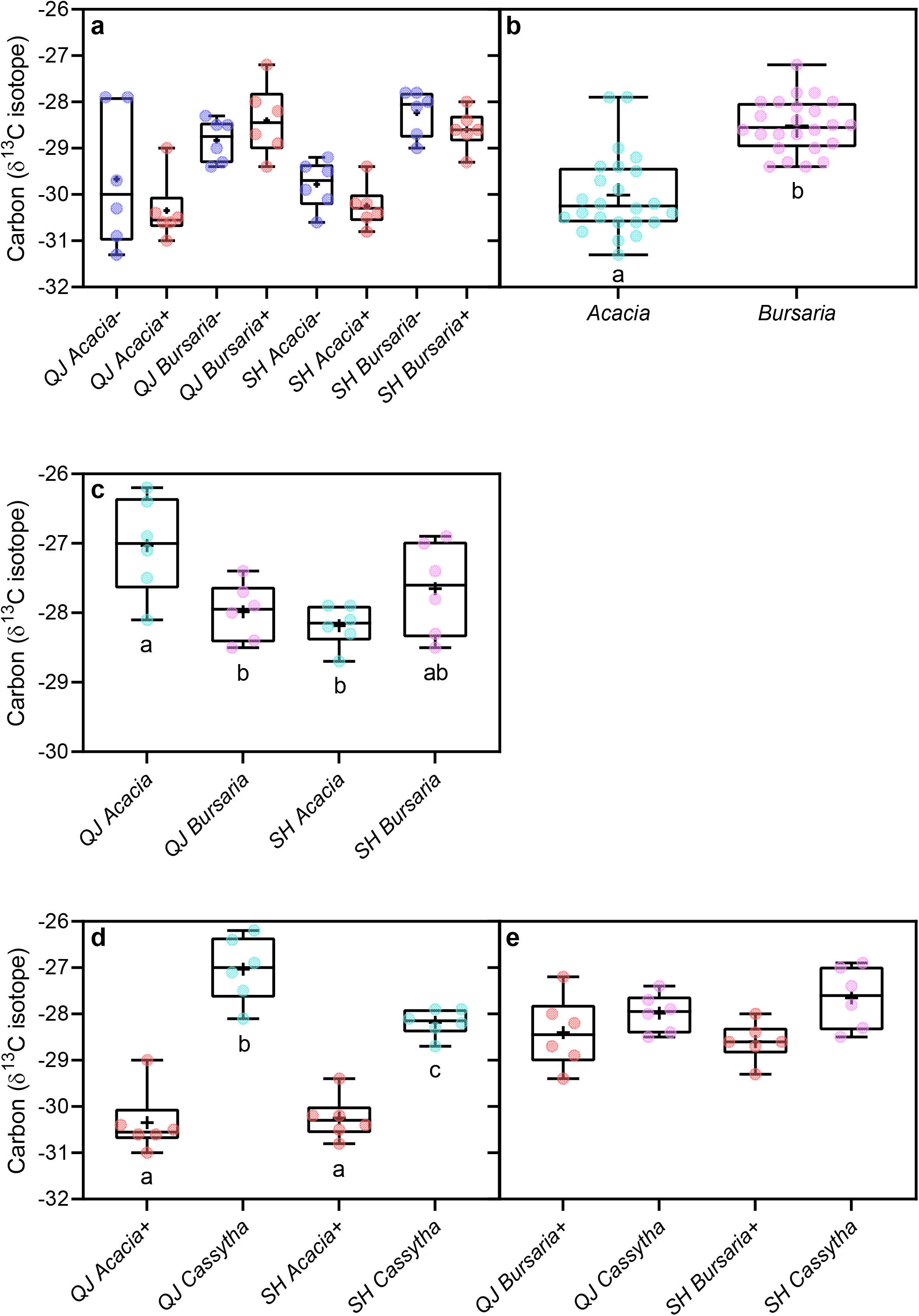
(**a**) δ ^13^C of *Acacia pycnantha* and *Bursaria spinosa* either uninfected (–) or infected (+) with *Cassytha pubescens* at two sites: Queens Jubilee (QJ) or Saddle Hill (SH). (**b**) Main effect of host species on host δ^13^C. (**c**) Parasite stem δ^13^C when infecting *A. pycnantha* or *B. spinosa* at QJ or SH. (**d**) δ^13^C of infected *A. pycnantha* and associated parasite, and (**e**) δ^13^C of infected *B. spinosa* and associated parasite at QJ or SH. All data points, median, 1^st^ and 3^rd^ quartiles, interquartile range and mean (+ within box) are shown, different letters signify significant differences: *n* = 6 (**a**, **c**, **d**, **e**) and *n* = 24 (**b**)

## Discussion

Our hypothesis that *C. pubescens* would negatively affect *R. anglocandicans* at both sites was confirmed, with significant effects of infection observed on a range of host physiological traits. *Cassytha pubescens* strongly affected host photosynthetic performance and stomatal conductance, regardless of site differences. Our second prediction that *C. pubescens* would have less impact on the native hosts, *A. pycnantha* and *B. spinosa*, was also mainly supported; although there was a significant negative impact on host stomatal conductance.

The negative effect of *C. pubescens* on *F*_v_/*F*_m_ and Φ_PSII_ of invasive *R. anglocandicans* across sites has also been found for other invasive hosts (*U. europaeus* and *Cytisus scoparius*) infected with this native parasite (Shen et al. 2010; Cirocco et al. 2018, 2021a; but see Prider et al. 2009). Similarly, Shen et al. (2013) found that *Cuscuta campestris* had a strong negative effect on apparent quantum yield of the major invasive host, *Mikania micrantha*. In contrast, *C. pubescens* generally has no effect on *F*_v_/*F*_m_ and Φ_PSII_ of the native hosts, *Leptospermum myrsinoides* or *Acacia paradoxa* (Prider et al. 2009; Cirocco et al. 2015, 2021b, but see the latter study). This is becoming a consistent finding for native hosts as we also found that *C. pubescens* had no effect on *F*_v_/*F*_m_ and Φ_PSII_ of *A. pycanatha* or *B. spinosa*. With regard to invasive host’s light use efficiency being negatively affected by *Cassytha* and *Cuscuta*, these findings may be due to negative parasite effects on host photosynthesis which was found in our study and Shen et al. (2013). As mentioned, declines in photosynthesis at a constant or increasing PFD would result in plants being exposed to excess light and if prolonged this can result in sustained photoinhibition. With regard to native hosts, their ETR was unaffected by the parasite which helps explain why their light-use efficiency was not impacted by *C. pubescens*. Another factor to consider is that native hosts may also have sufficient photoprotective capacity and xanthophyll engagement to cope with any excess light resulting from infection (Cirocco et al. 2015), whether invasive hosts have lower xanthophyll capacity requires investigation.

The negative parasite effect on ETR of *R. anglocandicans* has also been found for the invasive hosts *U. europaeus* and *Cytisus scoparius* (Prider et al. 2009; Shen et al. 2010; Cirocco et al. 2016b, 2017, 2018, 2020, 2021a). Similarly, a native parasite in China (*Cuscuta australis*) also negatively affected photosynthesis of the invasive hosts *Mikania micrantha* (Le et al. 2015) and young *Bidens pilosa* (Li et al. 2015). In the current study ETRs of the native hosts *A. pycnantha* and *B. spinosa* were unaffected by the parasite. Reports of the effect of *C. pubescens* on photosynthesis of native hosts are mixed. For example, the parasite has been found to negatively affect photosynthesis (but typically not growth) of the native host *Leptospermum myrsinoides* (Prider et al. 2009; Cirocco et al. 2016a). On the other hand, Cirocco et al. (2017, 2021b) found that *C. pubescens* had no effect on photosynthesis of *A. paradoxa*. Studies in Africa have found that the native root hemiparasite *Striga hermonthica* affects photosynthesis of native grasses in some cases (Gurney et al. 2002), but not others (Watling and Press 1998). *Striga* spp. have major effects on cereal crops in Africa, and are a component of natural ecosystems where native grasses may have evolved resistance/tolerance to infection (Gurney et al. 2002). Here, the effect of *C. pubescens* on ETR of *R. anglocandidans* may be due to effects on stomatal conductance (Fig. 2b), considering host N was not affected by infection. On the other hand, the lack of an infection effect on ETR of *A. pycnantha* and *B. spinosa* may be due to the negative effect of *C. pubescens* on host stomatal conductance not being large enough to affect photosynthesis of these native hosts.

Parasitic plants are commonly found to negatively affect host stomatal conductance, as found here for both *R. anglocandidans* and the two native hosts. *Cassytha pubescens* was also found to negatively affect g_s_ of the invasive *Cytisus scoparius* (Shen et al. 2010) and that of the native host *L. myrsinoides* (Cirocco et al. 2016a). A study in China also found that the native holoparasite *Cuscuta australis* negatively affected g_s_ of the invasive *M. micrantha*, especially in low water conditions (Le et al. 2015). Shen et al. (2007, 2013) also found that *Cuscuta campestris* strongly inhibited g_s_ of *M. micrantha*. Typically, the negative effects of infection on host g_s_ as in our study may be a response to water removal by the parasite.

δ^13^C of *C. pubescens* was significantly higher than that of *R. anglocandicans* at Horsnell but not at Morialta. This finding indicates that the parasite was more conservative in its water-use than the host at Horsnell, perhaps because hosts at this site were less hydrated as inferred from their significantly lower g_s_ as compared with plants at Morialta (Fig. 2c). *Cassytha pubescens* having significantly higher δ^13^C than invasive hosts has been reported previously (Cirocco et al. 2016b, 2018, 2021a). In the current study, we also observed that parasite δ^13^C was significantly higher than that of the two native hosts, *A. pycnantha* and *B. spinosa* as similarly found for the *C. pubescens-A. paradoxa* association (Cirocco et al. 2021b). This suggests, counter to observations of other hemiparasitic plants, that *C. pubescens* is more conservative in its water-use than its hosts, particularly in low water conditions. We have observed in glasshouse experiments that when the host is experiencing water deficits, *C. pubescens* wilts before the host, suggesting that even with stomatal closure the parasite may not be able to avoid desiccation and turgor loss. In contrast, mistletoes have high leaf tissue capacitance which may help explain why they are less conservative in their water-use (lower δ^13^C) than their hosts (Glatzel 1983; Davidson et al. 1989; Scalon and Wright 2015).

Another factor affecting parasite δ^13^C may relate to nitrogen availability. Schulze et al. (1984) hypothesised that xylem-tapping parasites maintain high transpiration rates in order to acquire sufficient N from their hosts. Moreover, the parasite may become more conservative in its water-use (e.g. less negative δ^13^C) when its N requirements are met which may be expedited when the host is rich in N. There is evidence that mistletoes have less negative δ^13^C when attached to legume (N-rich) versus non-legume hosts (Schulze and Ehleringer 1984; Ehleringer et al. 1985; but see Scalon and Wright 2015). We also found support for this as the difference in δ^13^C between *C. pubescens* and its host, was greater when infecting the leguminous host *A. pycnantha* than the non-legume *B. spinosa* (Fig. 8d, e). In accordance with *A. pycnantha* having significantly higher foliar [N] than *B. spinosa* (Fig. 7c), the parasite also had significantly higher [N] when infecting the former host (Fig. 7e). Similarly, nitrogen concentration of *C. pubescens* has also reflected that of both native and invasive hosts in other studies (Cirocco et al. 2016b, 2020, 2021a, b). For the *C. pubescens-R. anglocandicans* association, parasite [N] was also similar to that of the invasive host. Scalon and Wright (2015) also found that for 168 mistletoe:host pairs, parasite N was strongly positively correlated with host N.

## Conclusions

The results clearly demonstrated that *C. pubescens* had a strong negative impact on physiology of the invasive *R. anglocandicans* across sites. This supports previous studies that have shown significant negative impacts on invasive hosts when infected by native parasites. In contrast, the parasite had no effect on photosynthesis or photoinhibition of the two native hosts; although g_s_ was negatively affected. Native hosts showing higher tolerance might be due to *C. pubescens* forming less effective haustorial connections to their vasculature, as has been found for another native host, *Acacia myrtifolia* (Facelli et al. 2020). Alternatively, the more conservative use of resources by these native sclerophyllous shrubs may limit the resources available to the parasite. Nevertheless, further work is required to evaluate the impact of this native parasite on its native hosts, including long-term monitoring of host growth and reproduction. It seems that, unlike other hemiparasites, *Cassytha* is unable to increase its uptake of host resources by increasing g_s_, as inferred from its higher δ^13^C than the host. In fact, when N becomes more available, it seems to lower g_s_ even further. This conservative parasite water-use may help explain why this generalist, native parasite is not found in more arid to semi-arid regions of Australia, unlike native mistletoes.

## Author contribution statement

RMC and JMF conceived and designed each of the studies. RMC performed the studies and analysed the data. RMC, JMF, and JRW interpreted the analysis and wrote the manuscript.

## Supplementary Information

The online version contains supplementary material available

**Fig. S1** Photos of *R. anglocandicans* uninfected or infected with *C. pubescens* at the two sites

**Fig. S2** Photos of *A. paradoxa* and *B. spinosa* uninfected or infected with *C. pubescens* at the two sites

**Fig. S3**–**S4** Light, air temperature and relative humidity when midday yield and stomatal conductance measurements were made

**Fig. S5** Photosynthetic performance of *C. pubescens* when infecting *R. anglocandicans*

**Fig. S6** Photosynthetic performance of *C. pubescens* when infecting *A. paradoxa* or *B. spinosa*

## Acknowledgements

Special thanks to Grace Porter-Dabrowski, Mitchell Groat and Bernardo J. O’Connor for their assistance with fieldwork, Dr Tony Hall (Mawson Analytical Spectrometry Services) and Jason Young (Flinders Analytical) for their expert mass spectrometry analysis. This work was supported by the Department of Agriculture, Water and the Environment (Australian Government) [55119480]; and the Department for Environment and Water (Adelaide and Mount Lofty Ranges Natural Resources Management Board) [55123913].

