## Supplementary material for "Impact of a native hemiparasitic plant on invasive and native hosts in the field": Supp Figs S1-S2

**Original Article**

**Horsnell Gully Conservation Park**

**a**

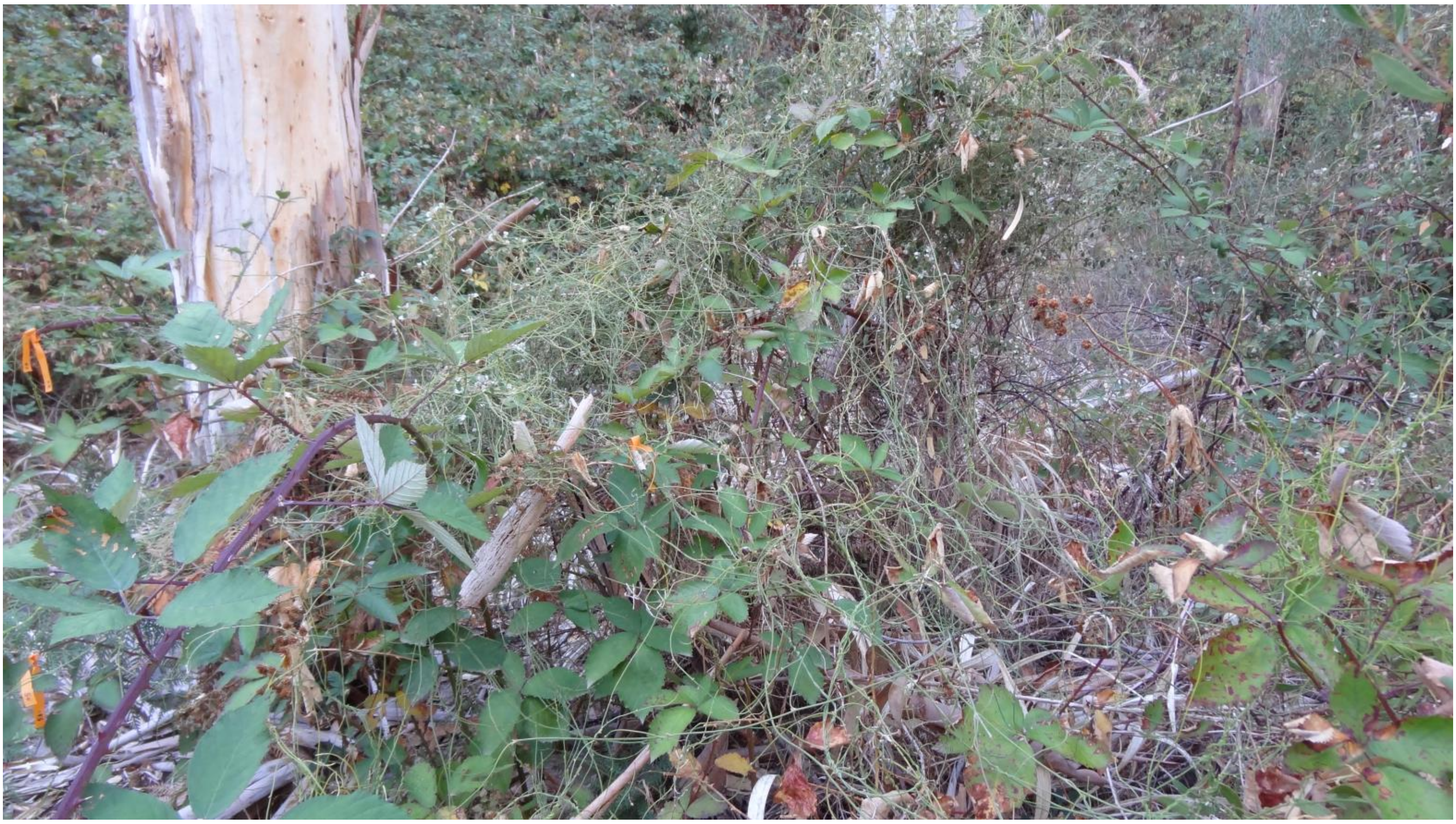

**b**

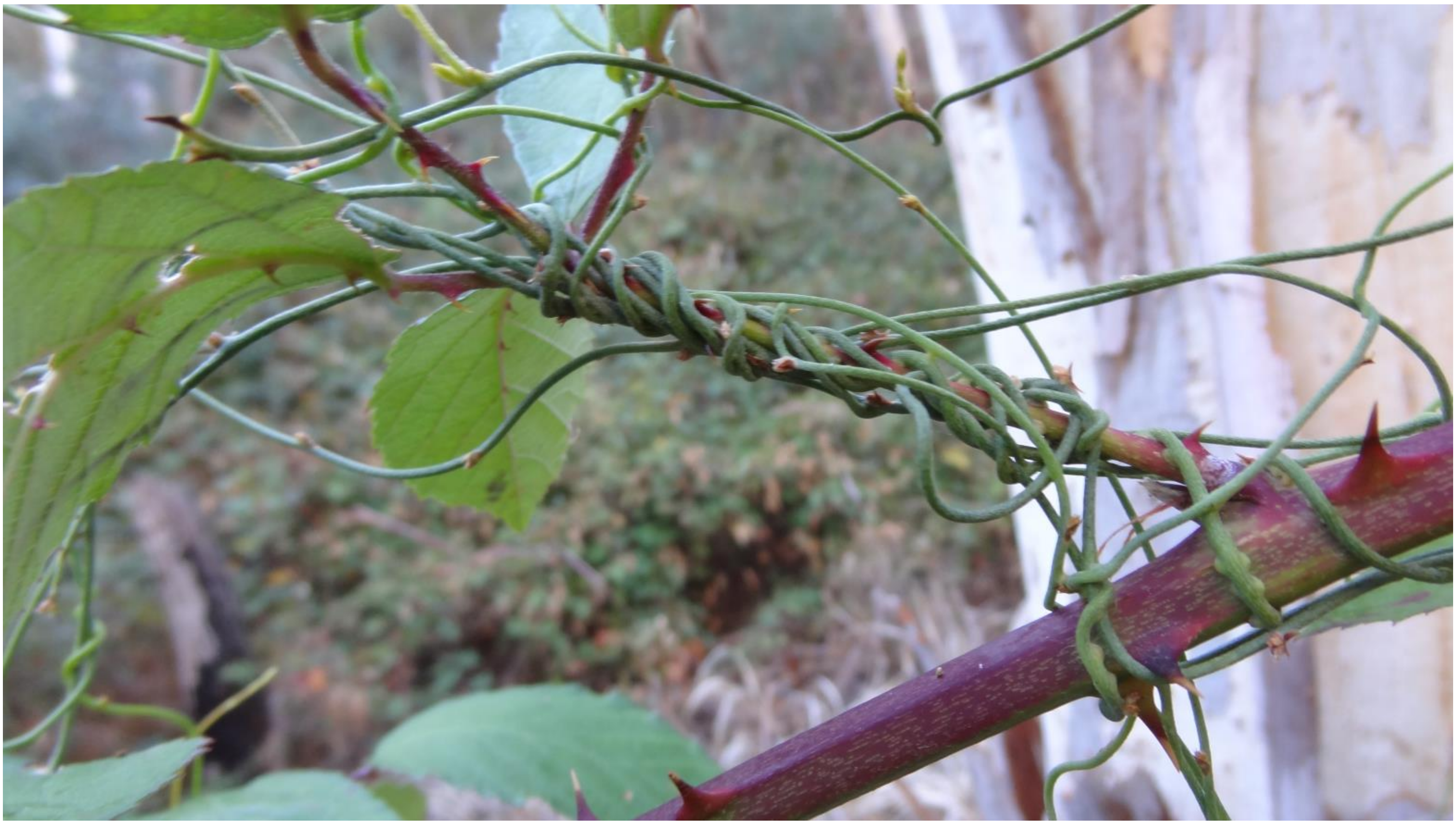

**Morialta Conservation Park**

**c**

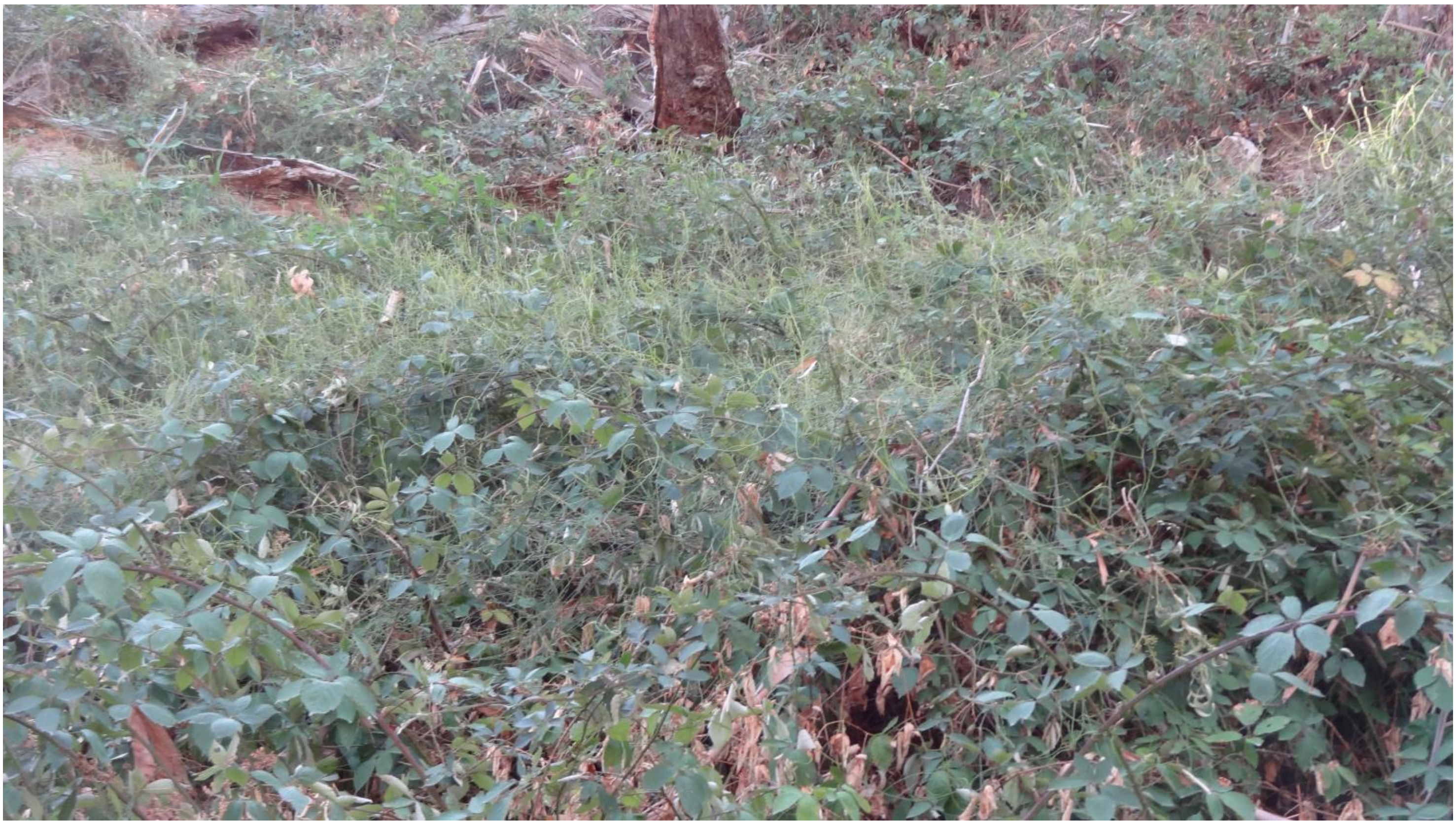

**d**

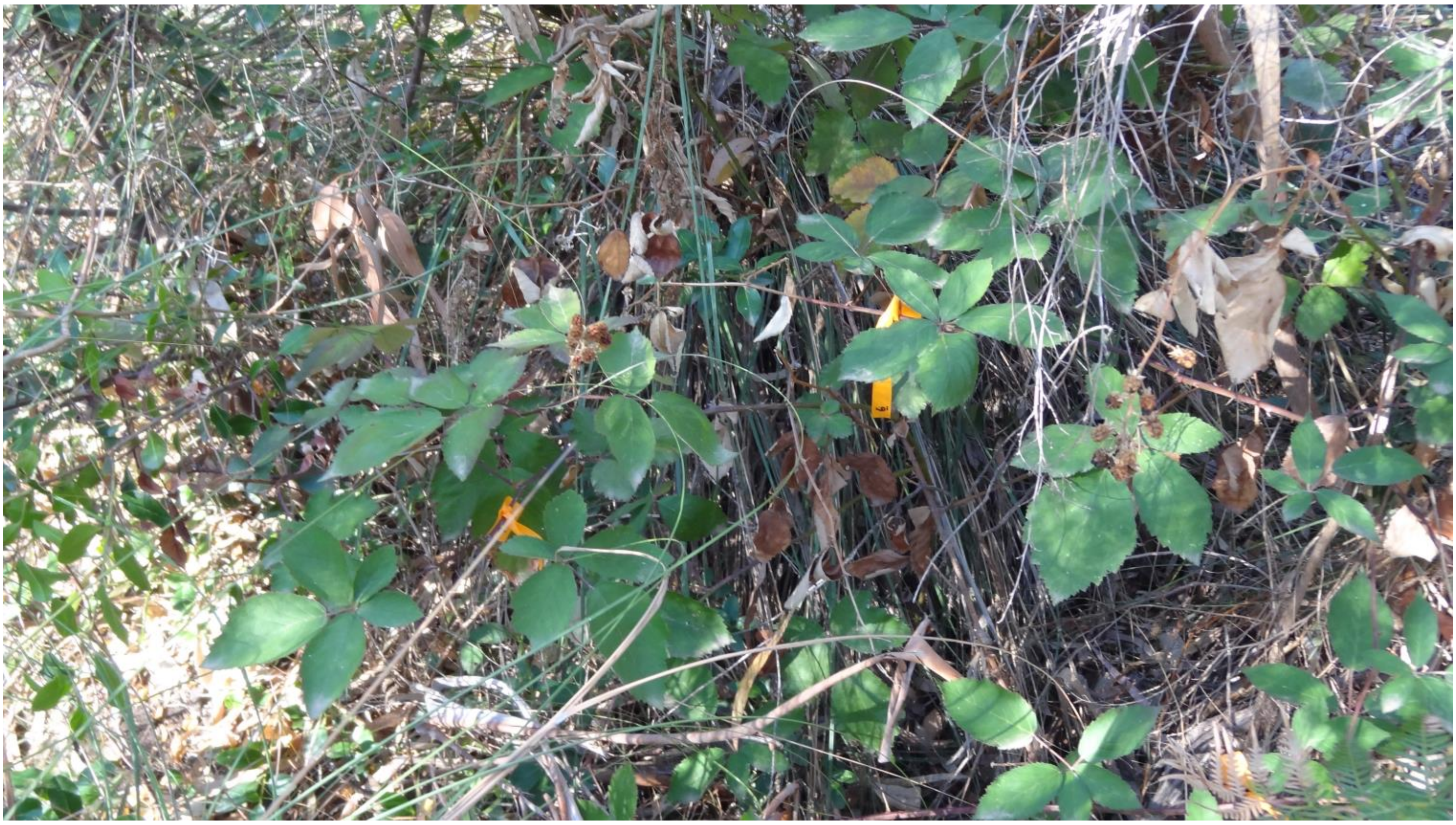

**e**

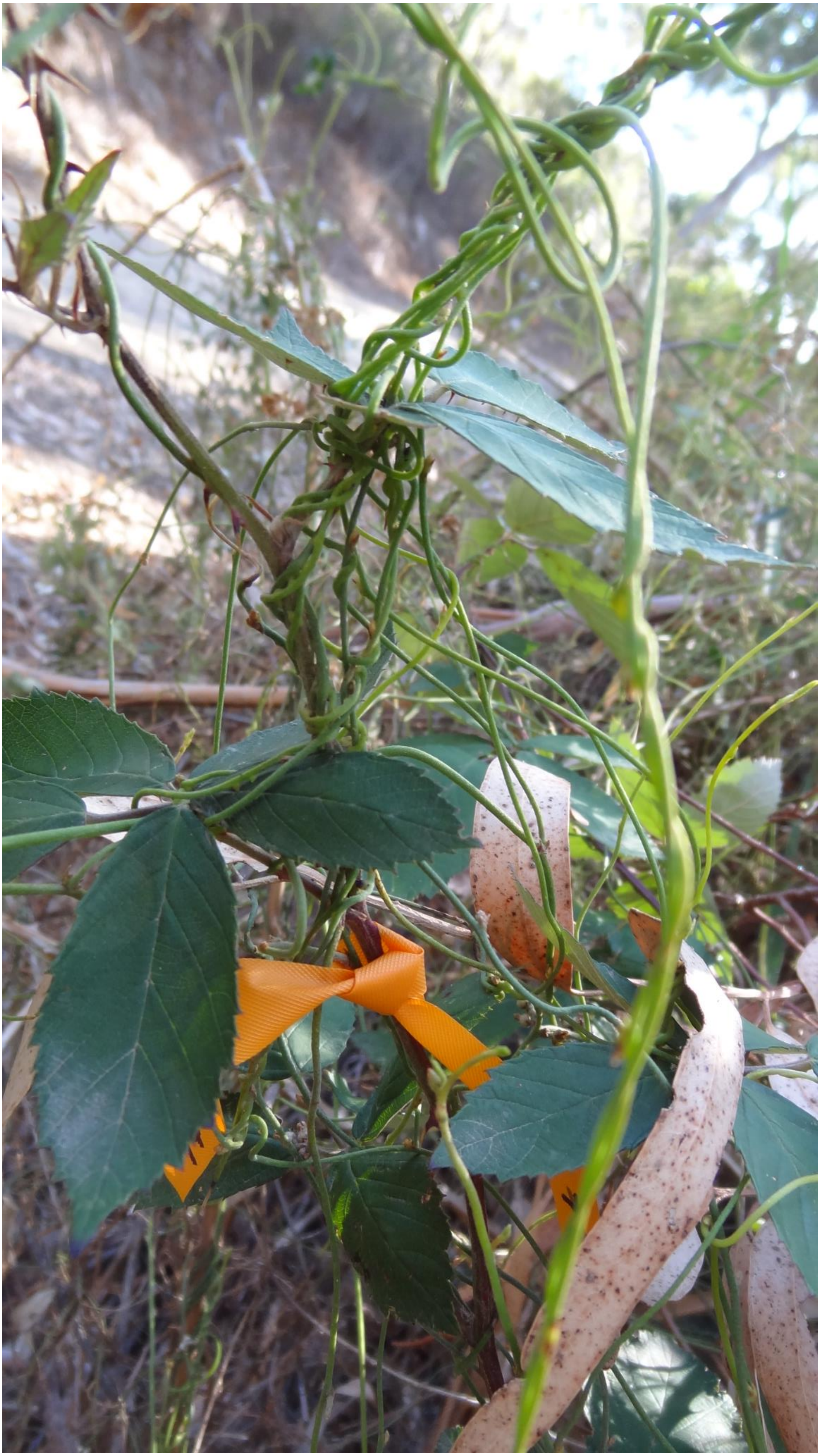

**f**

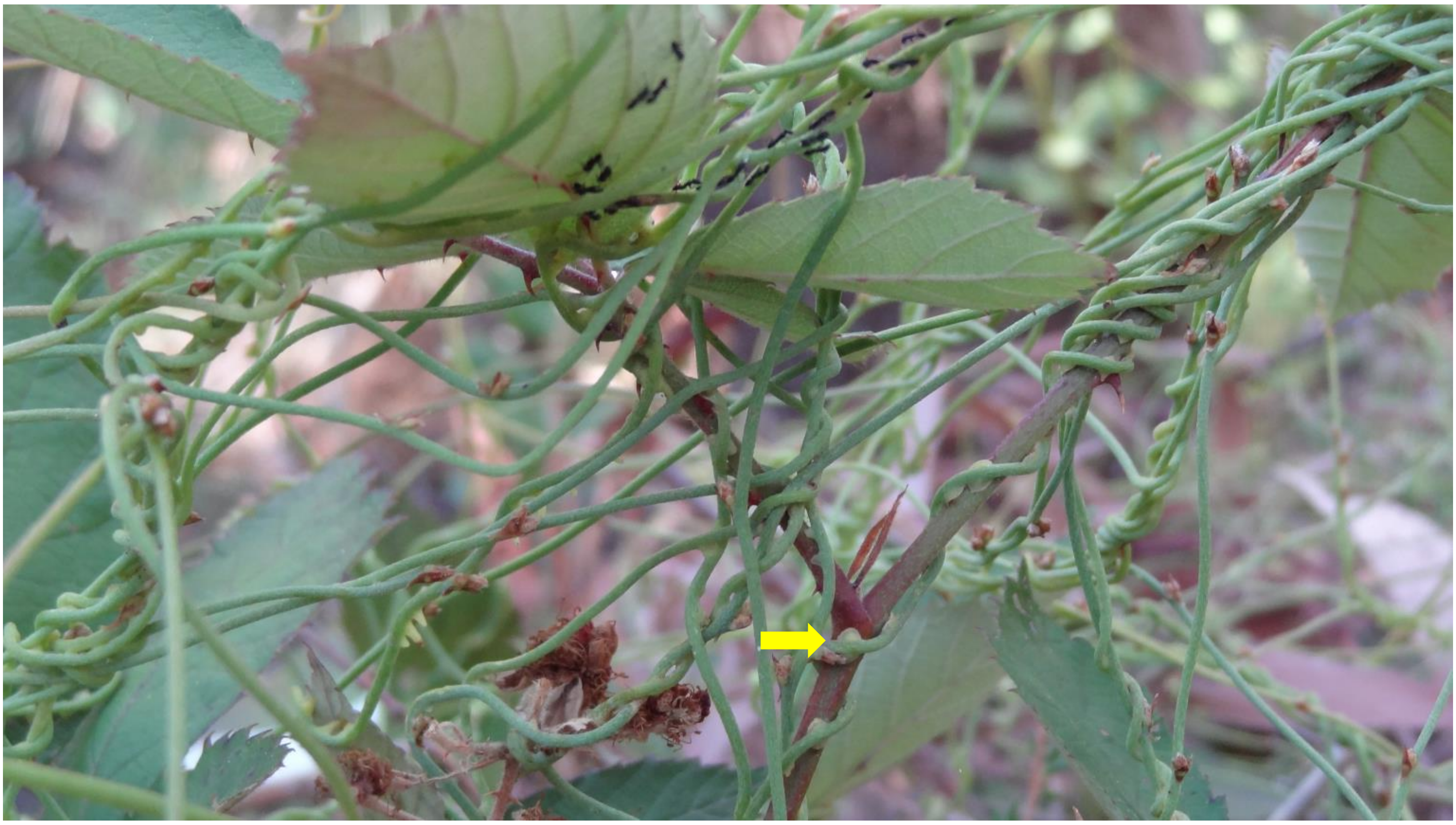

**Fig. S1** Photos of experimental plants (tagged with flagging tape) at the two field sites (Horsnell Gully and Morialta Conservation Parks), located in the Mt. Lofty Ranges of South Australia. Horsnell: (**a**) *Rubus anglocandicans* infected with *Cassytha pubescens* and (**b**) heavily infected host petiole. Morialta: (**c**) and (**d**) Host infected with parasite and uninfected plants used as controls, respectively. (**e**) Infected host and (**f**) infected petiole with yellow arrow indicating haustorial connection of *C. pubescens* to *R. anglocandicans*

**Saddle Hill Road**

**a**

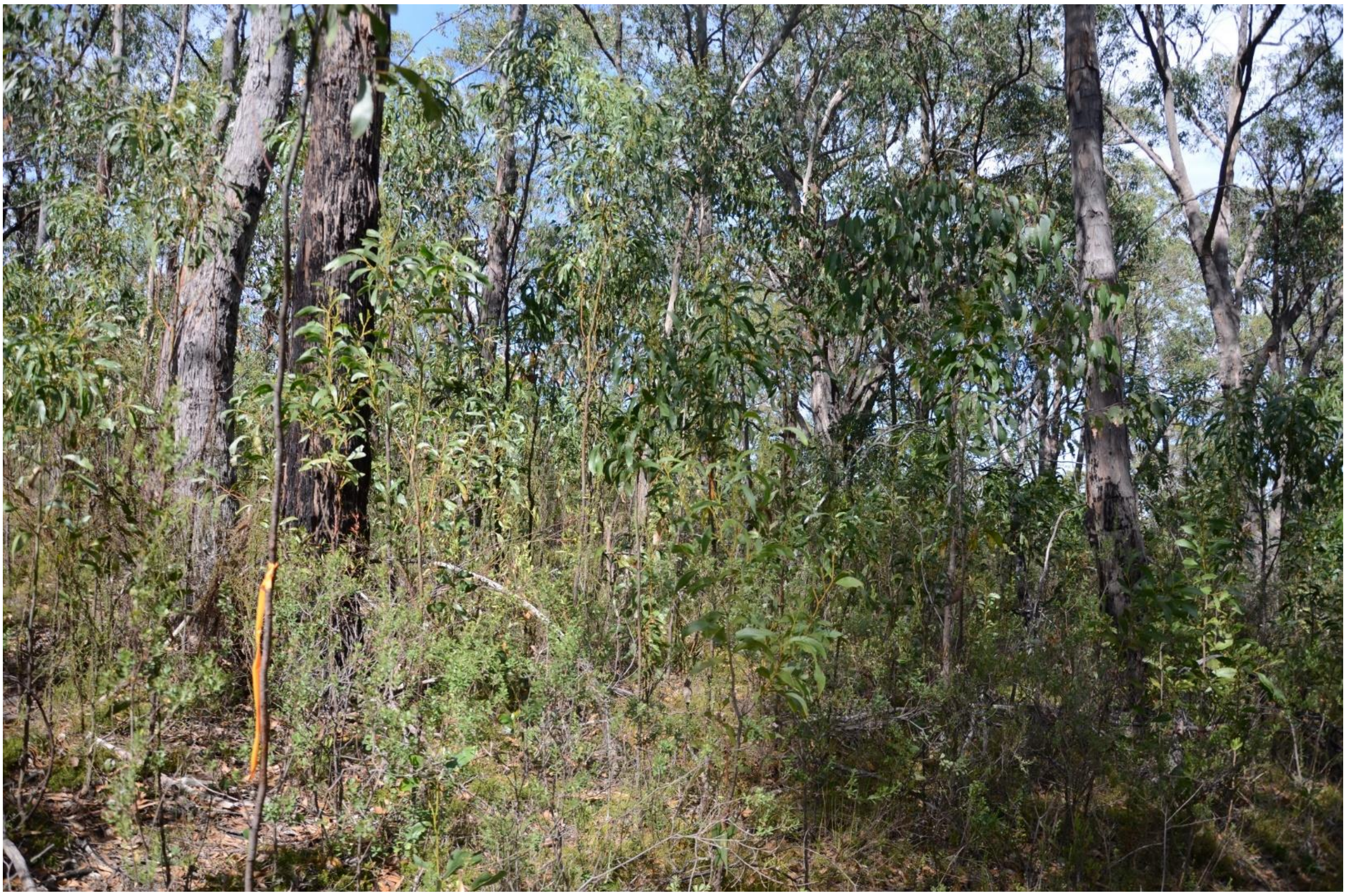

**b**

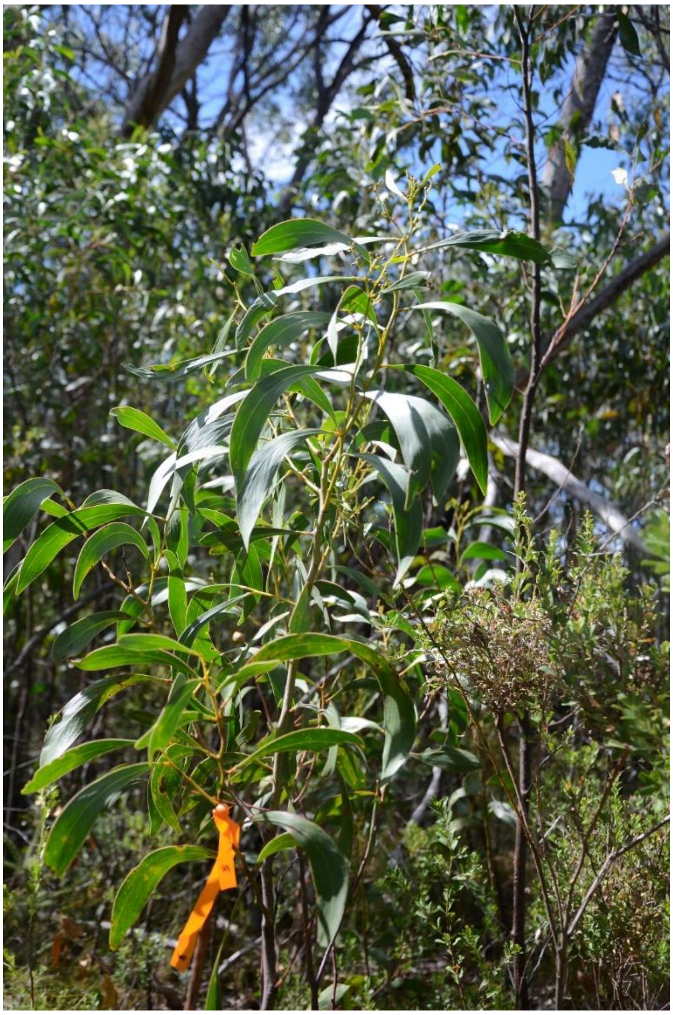

**c**

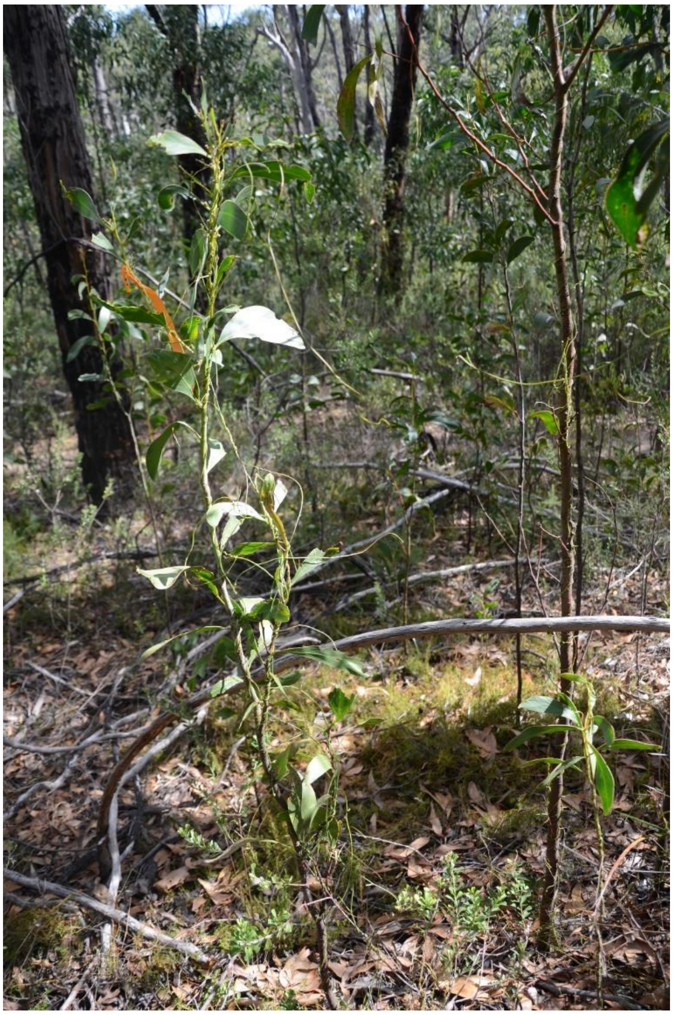

**d**

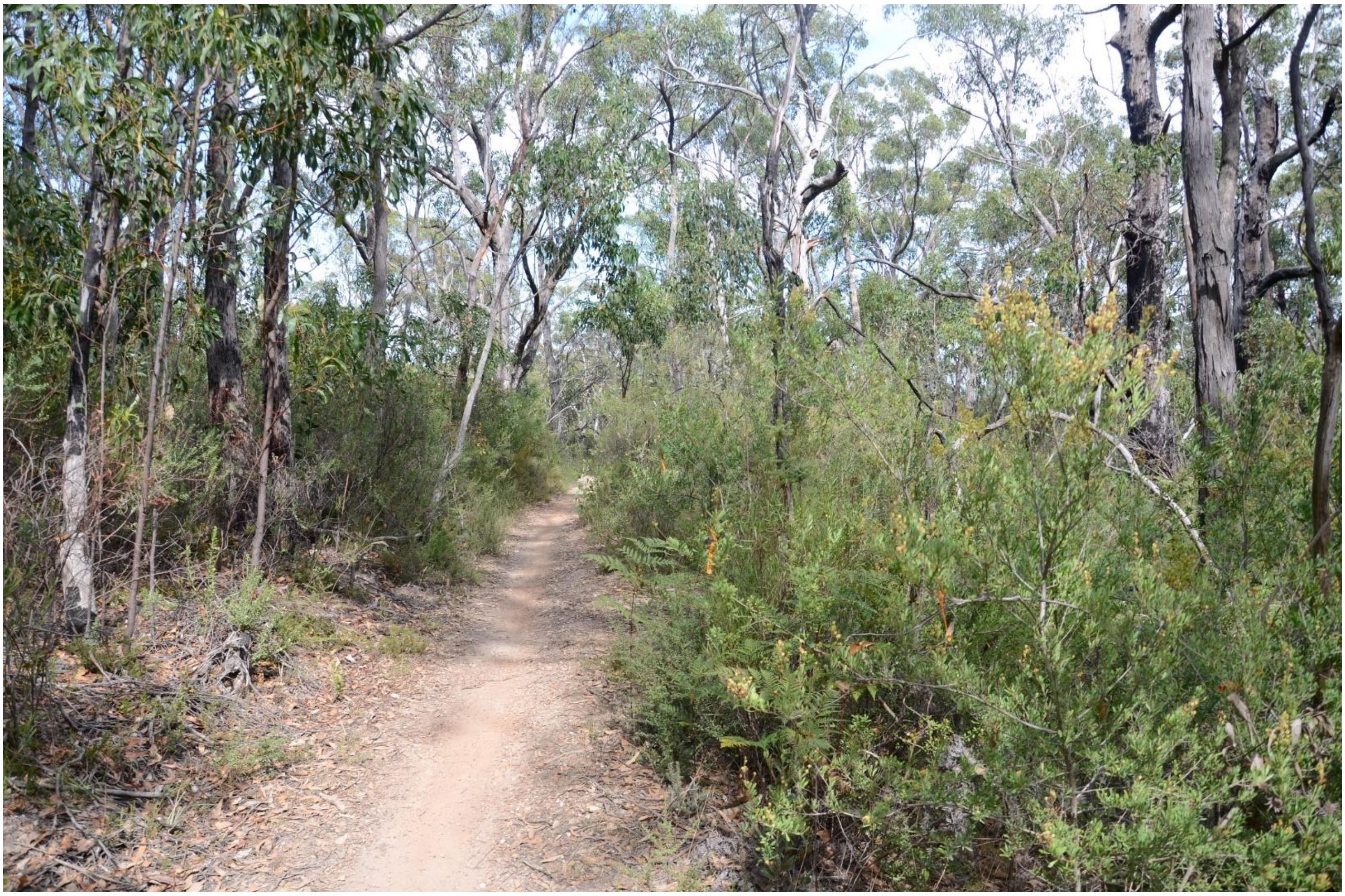

**e**

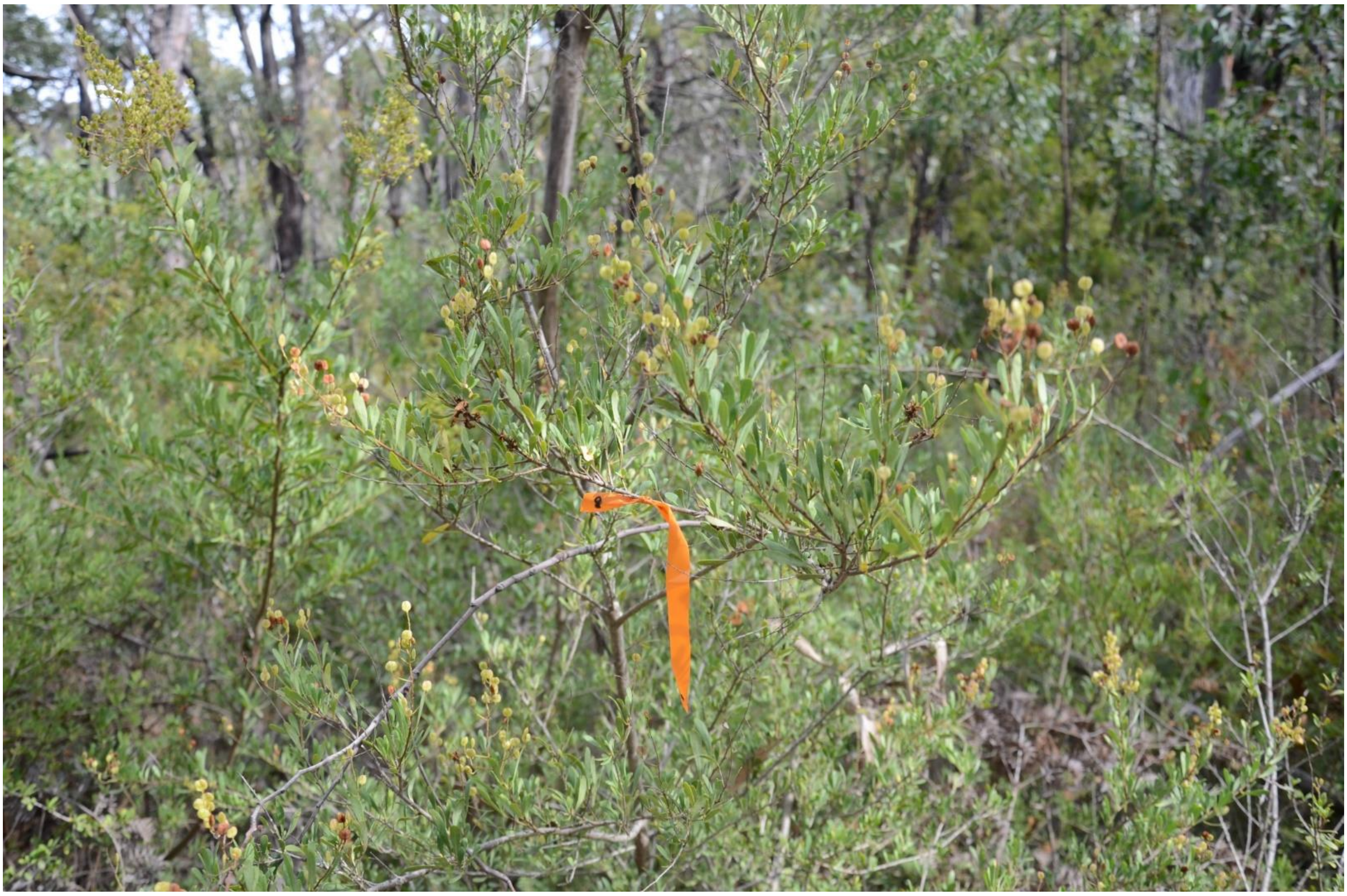

**f**

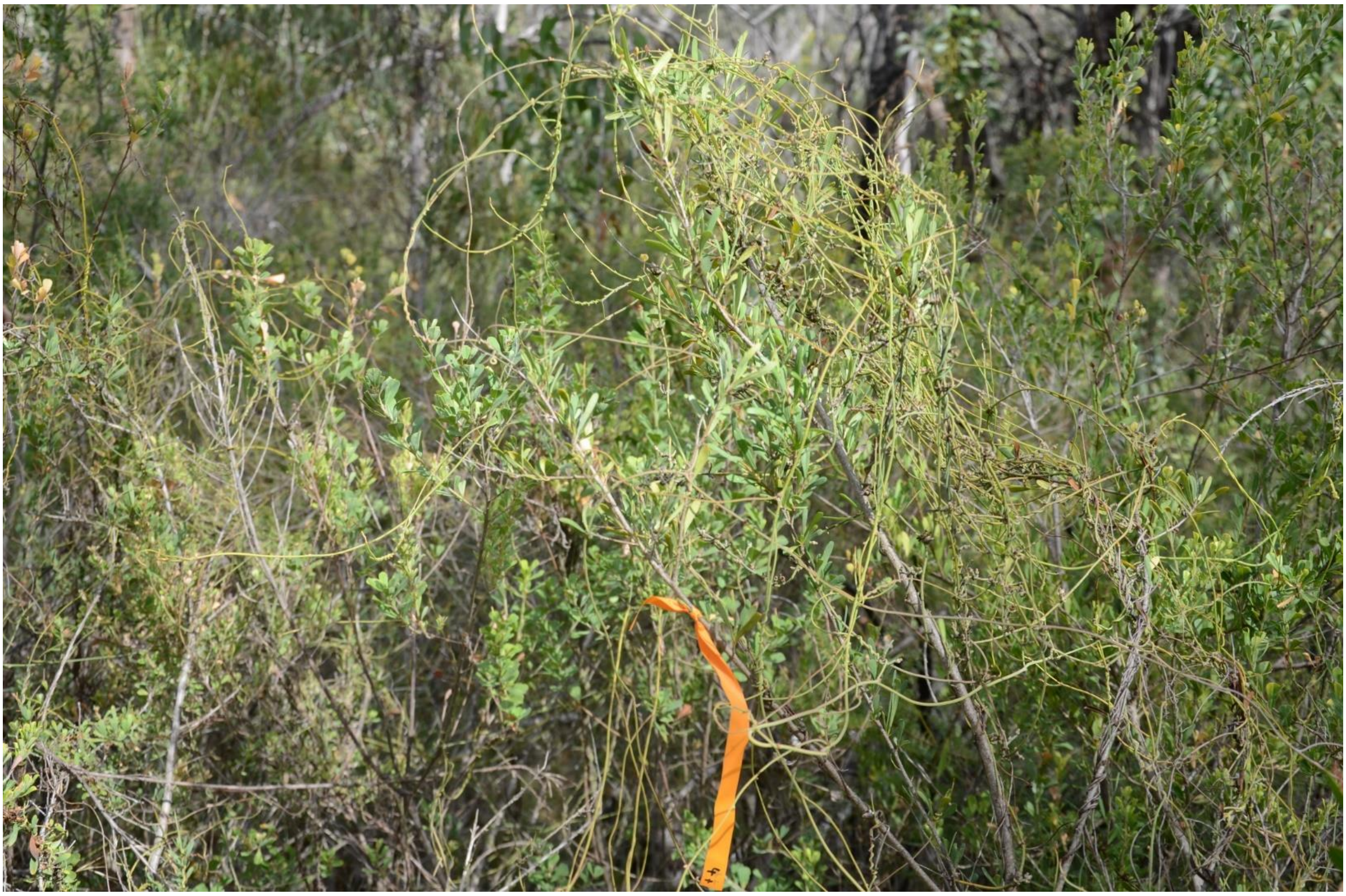

**Queens Jubilee Drive**

**g**

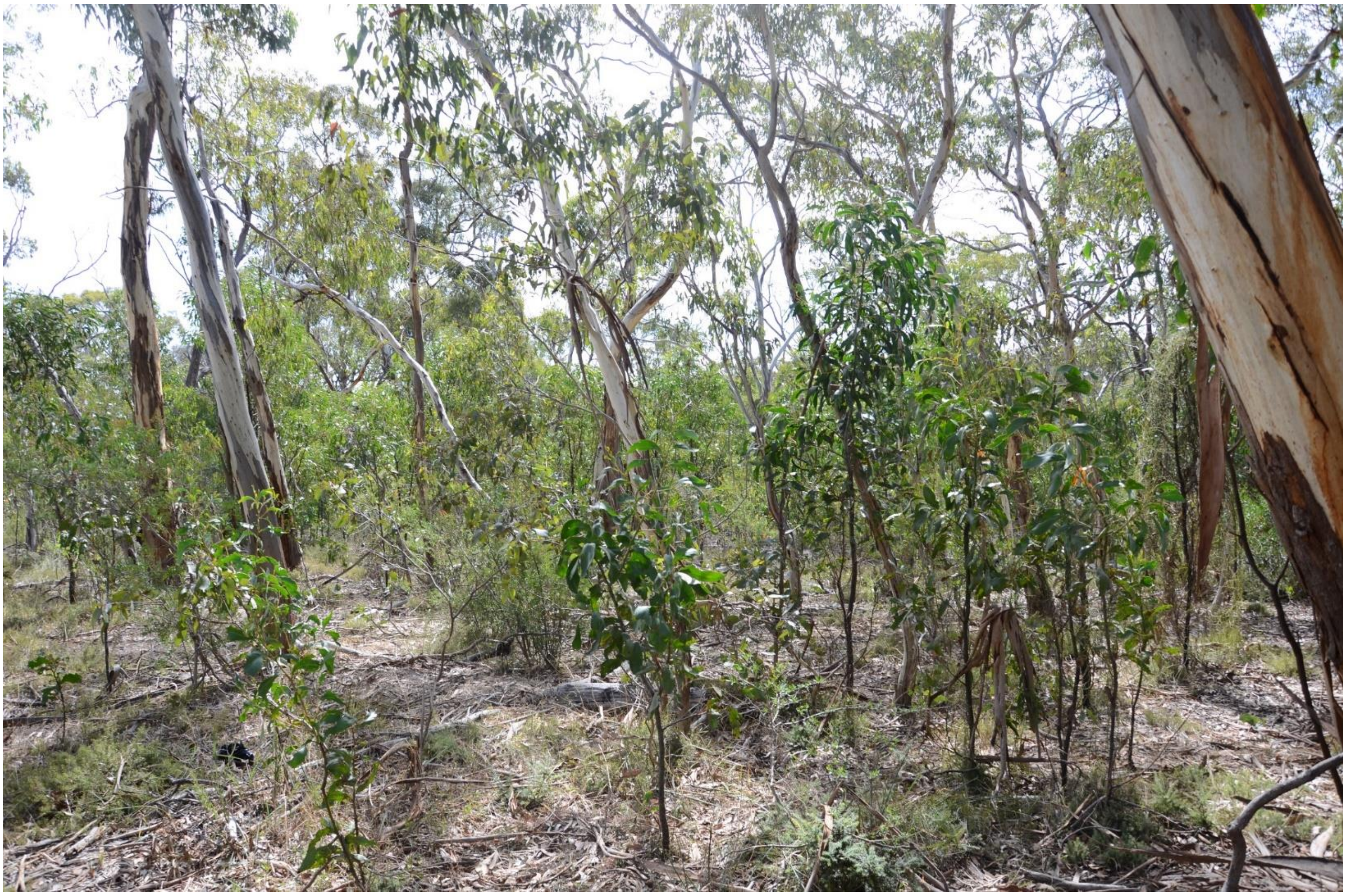

**h**

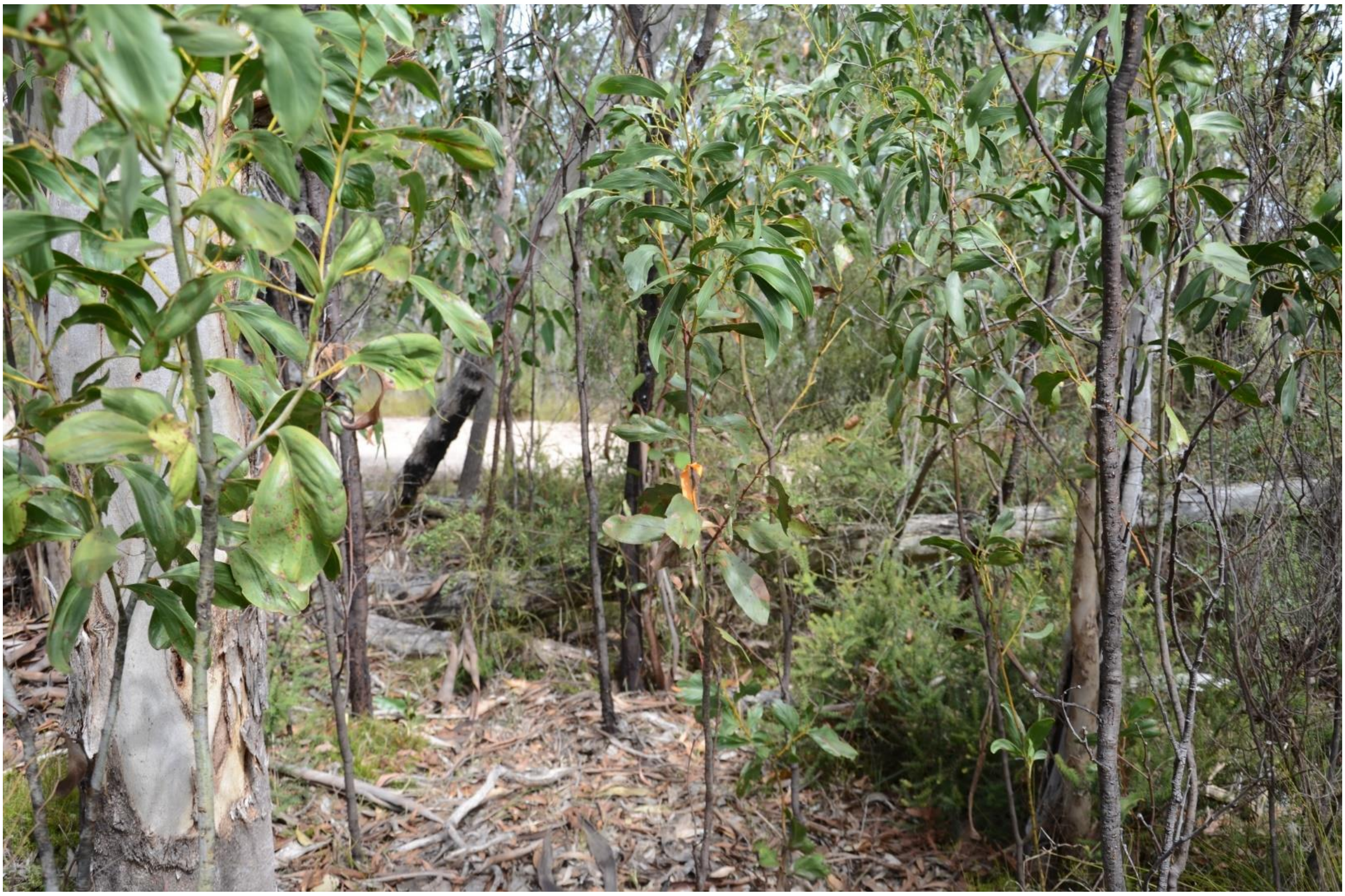

**i**

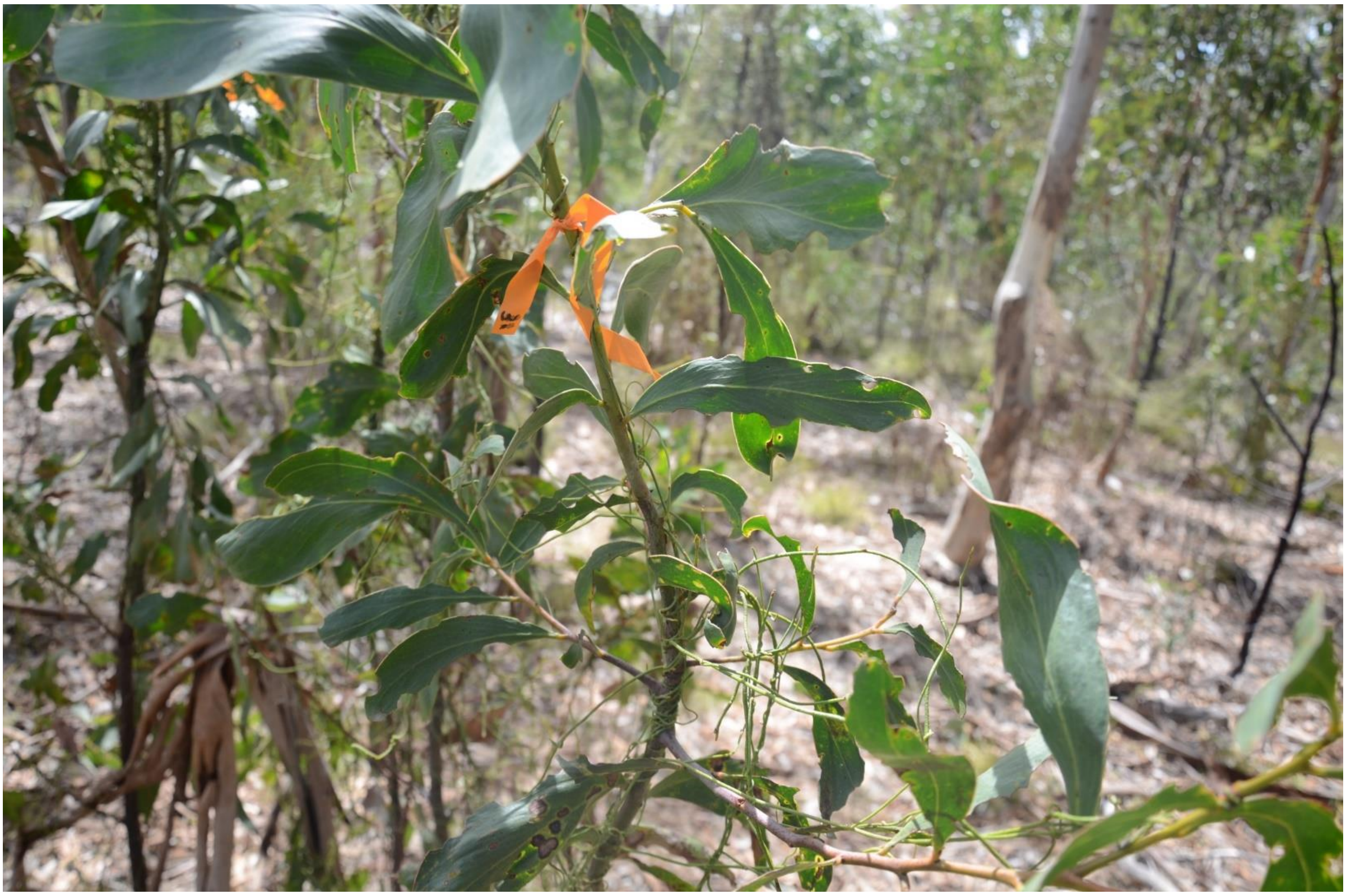

**j**

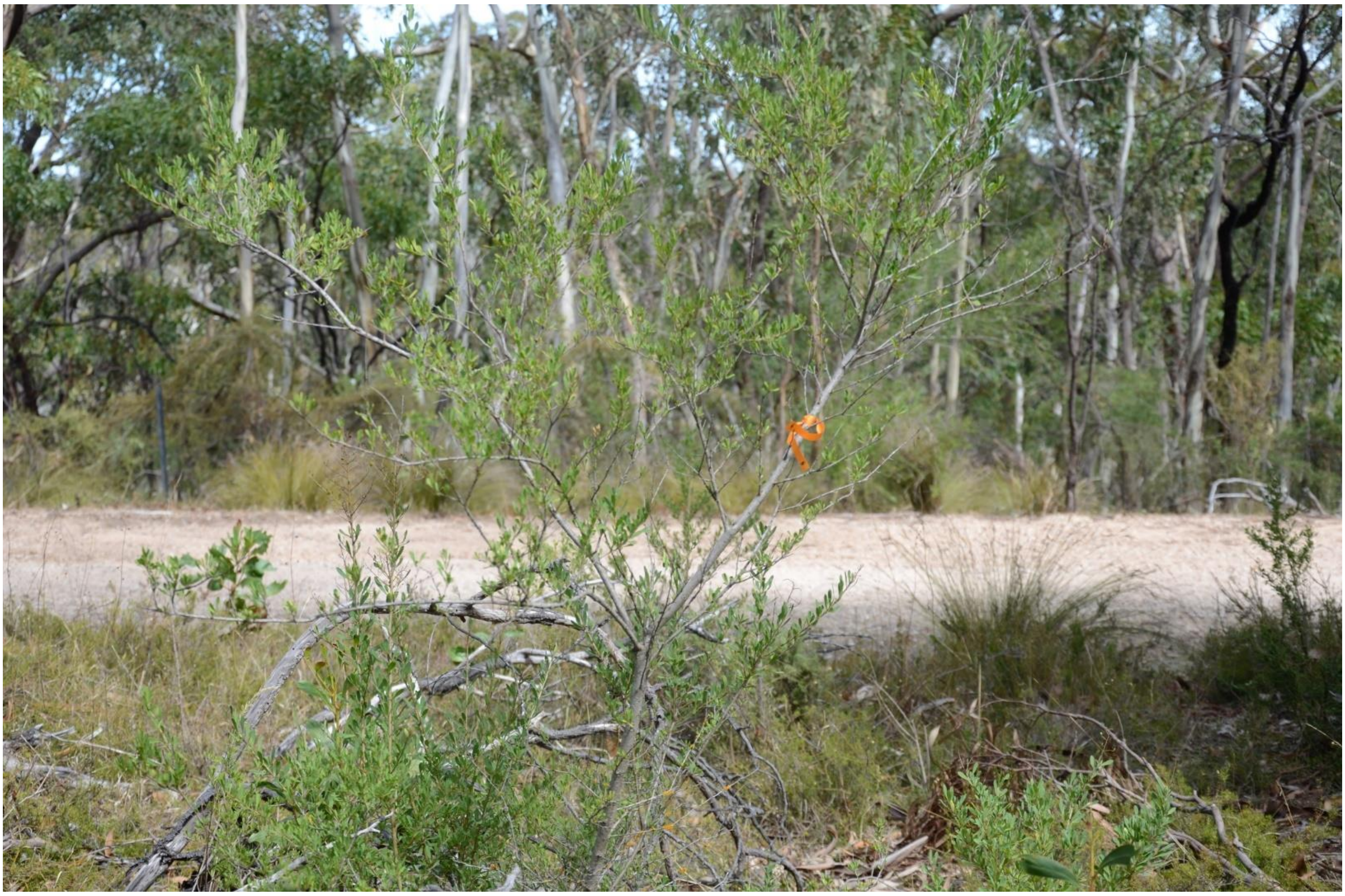

**k**

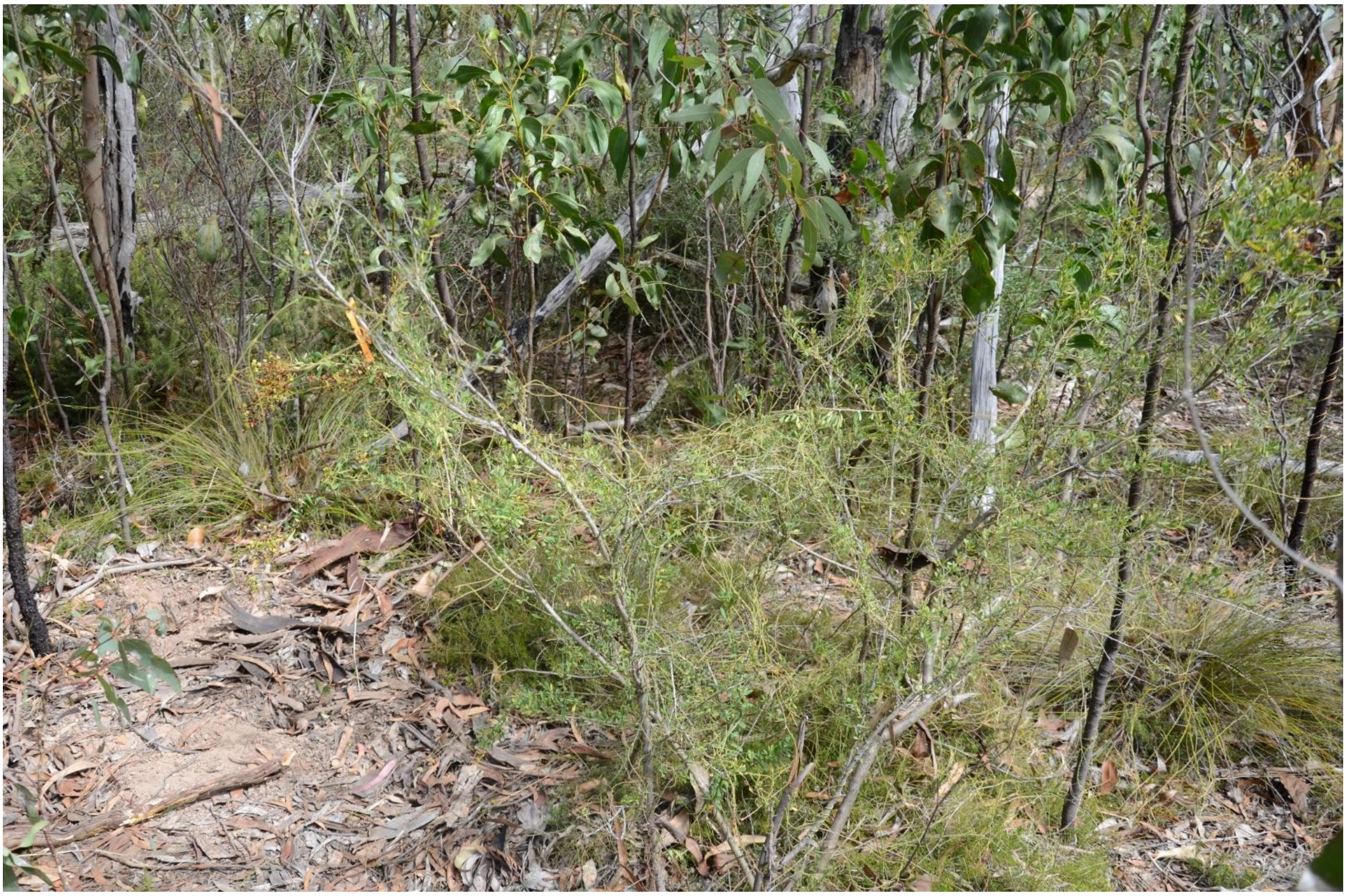

**l**

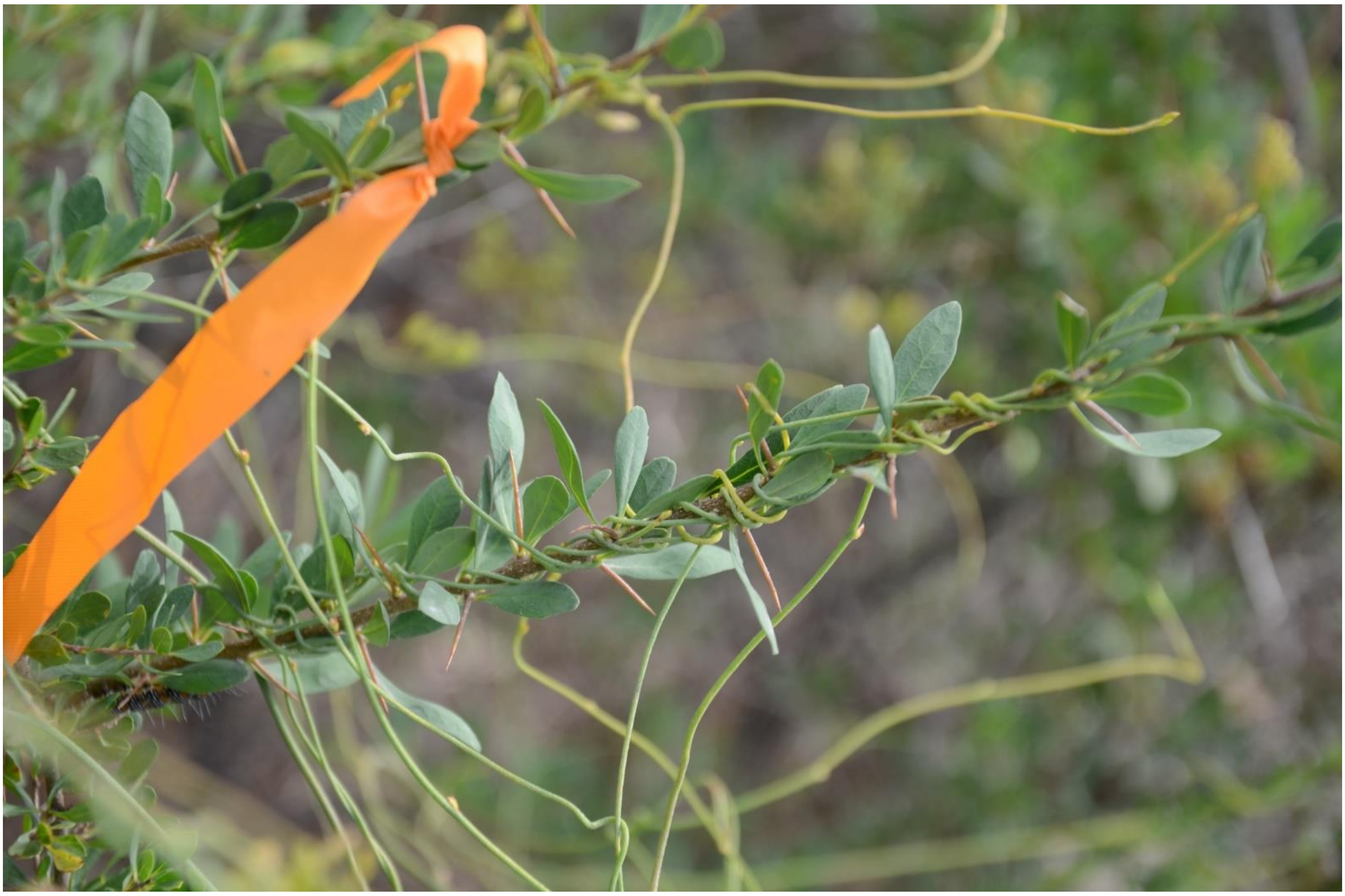

**m**

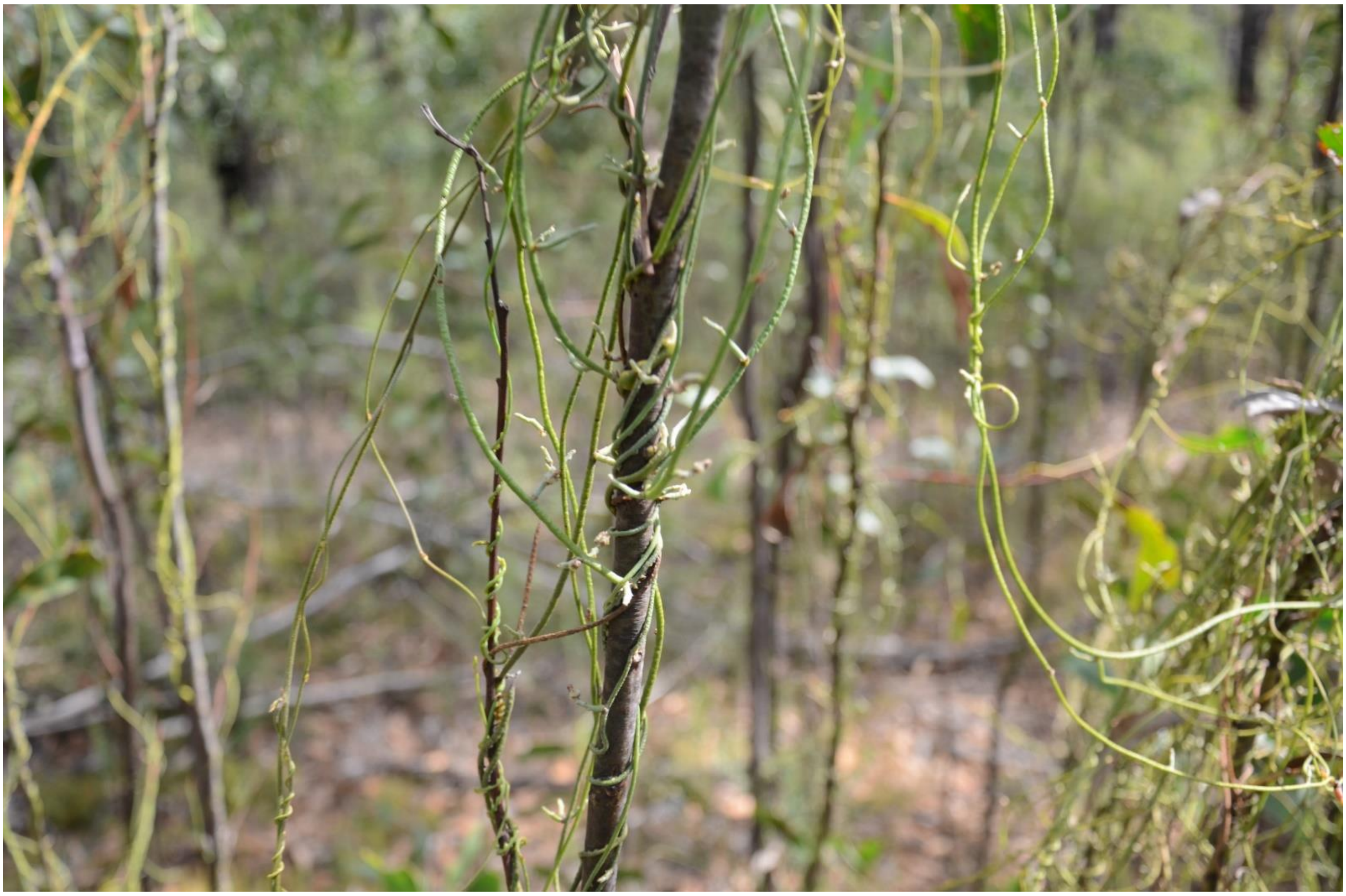

**Fig. S2** Photos of experimental plants (tagged with flagging tape) at the two field sites; Saddle Hill Rd: (**a**) general view of area comprising (**b**) uninfected and (**c**) *Cassytha* infected *Acacia pycnantha*. (**d**) general view of area comprising (**e**) uninfected and (**f**) *Cassytha* infected *Bursaria spinosa*. Queens Jubilee Drive: (**g**) general view, (**h**) uninfected and (**i**) *Cassytha* infected *A. pycnantha*, (**j**) uninfected and (**k**) *Cassytha* infected *B. spinosa*. Close-up of *Cassytha* infecting (**l**) *B. spinosa* and (**m**) *A. pycnantha*
