## Supplementary material for "Impact of a native hemiparasitic plant on invasive and native hosts in the field": Supp Figs S3-S6

**Original Article**

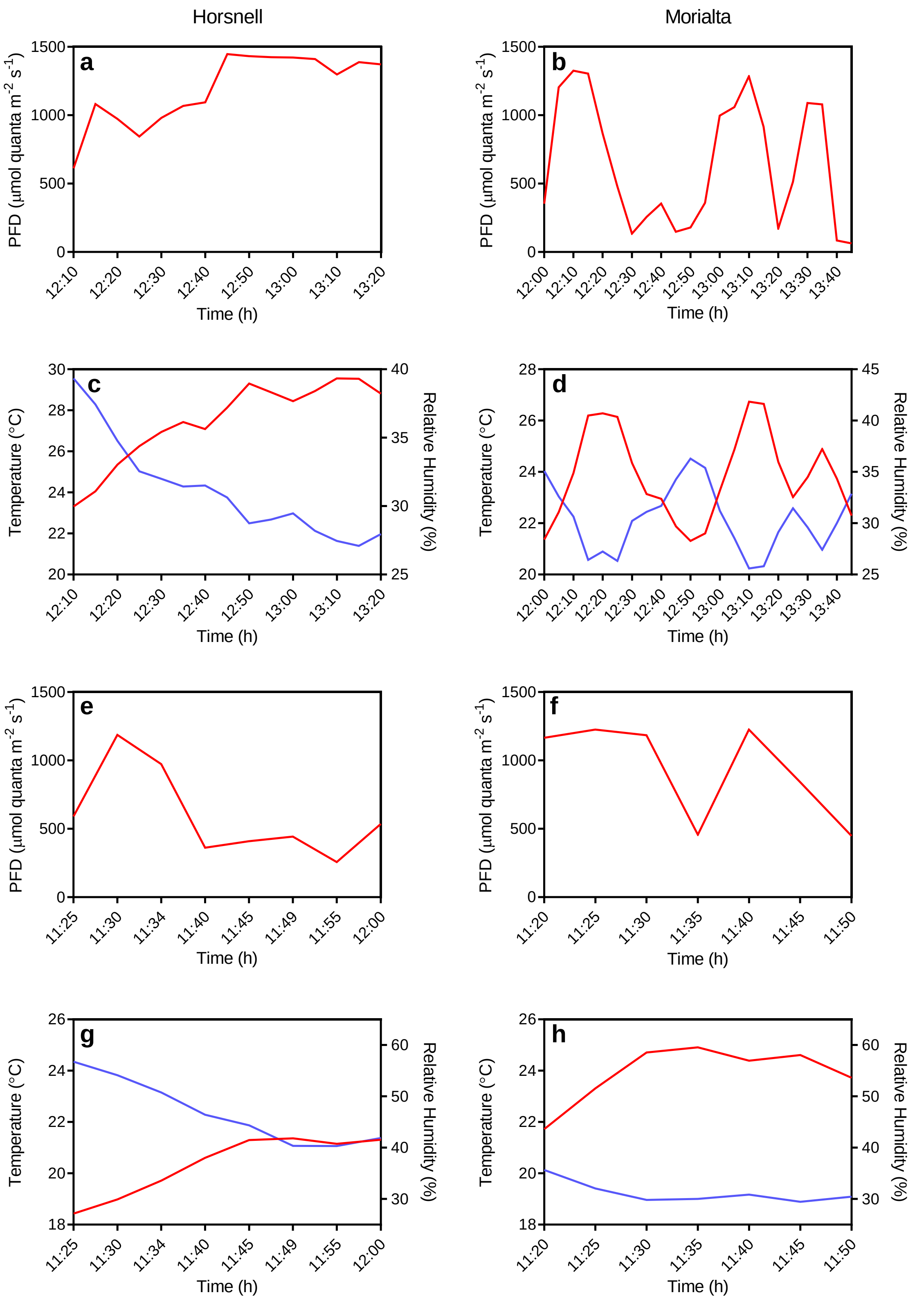


**Fig. S3** Top four graphs: (**a**) and (**b**) photon flux density and (**c**) and (**d**) air temperature (red, left axis) and relative humidity (blue, right axis), respectively, at two sites (Horsnell or Morialta) at the time/day when midday quantum yield ($\Phi_{PSII}$) electron transport rate (ETR)

measurements were made. Bottom four graphs: (**e**) and (**f**) photon flux density and (**g**) and (**h**) air temperature (red, left axis) and relative humidity (blue, right axis), respectively, at the two sites at the time/day when stomatal conductance measurements were made. $\Phi_{PSII}$ and ETR measurements were conducted on *Cassytha pubescens* and *Rubus anglocandicans* in mid-April 2019 and stomatal conductance measurements were made on the invasive host in late April 2019. Photosynthetic photon flux density (PPFD), air temperature and relative humidity were recorded with a LI-1400 data logger equipped with a quantum sensor (LI-190 SA) and relative humidity/air temperature sensor (1400-104) (LI-COR, Lincoln NEB.)

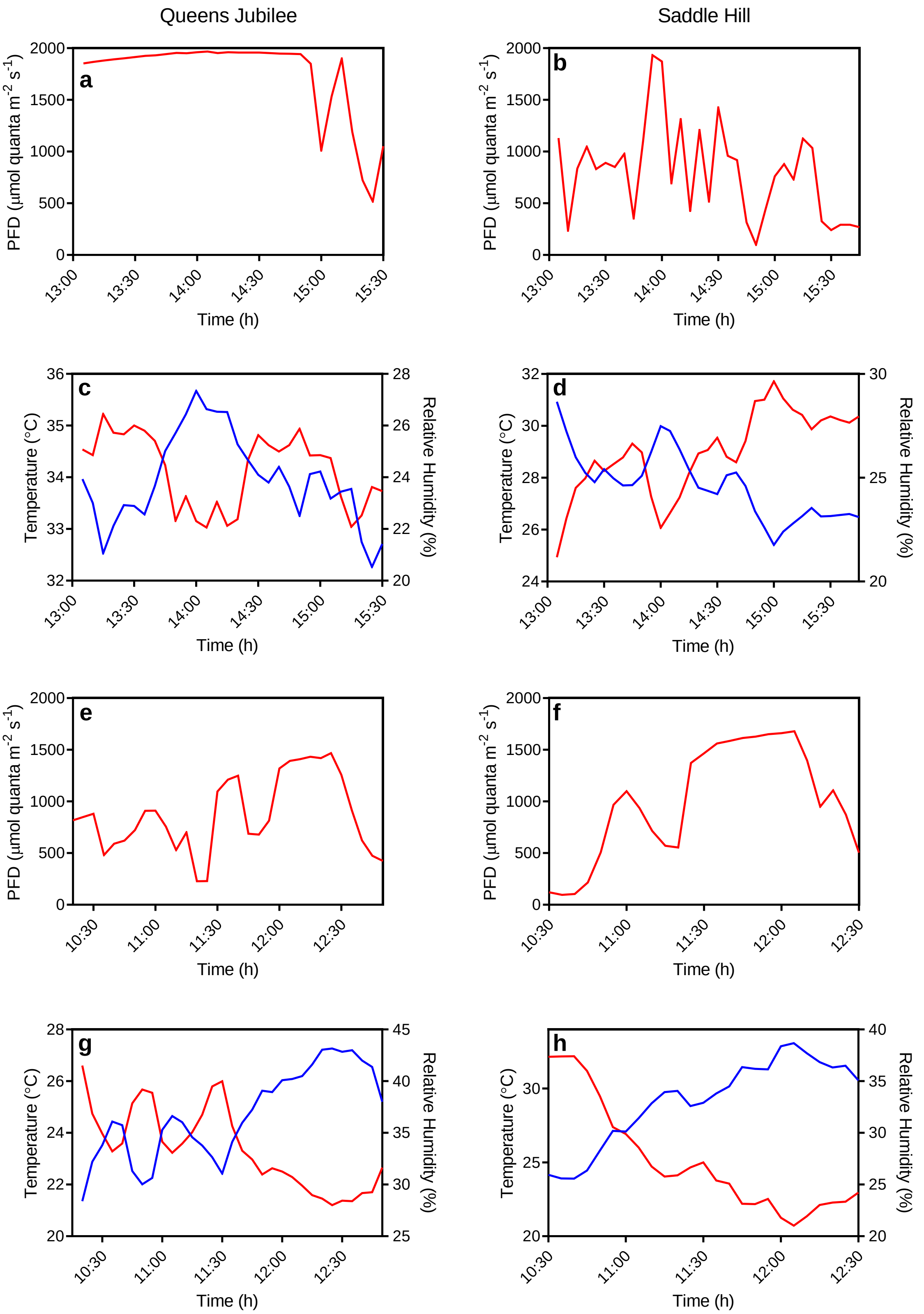


**Fig. S4**. Top four graphs: (**a**) and (**b**) photon flux density and (**c**) and (**d**) air temperature (red, left axis) and relative humidity (blue, right axis), respectively, at two sites (Queens Jubilee or Saddle Hill) at the time/day when midday quantum yield ($\Phi_{PSII}$) electron transport rate (ETR)

measurements were made. Bottom four graphs: (**e**) and (**f**) photon flux density and (**g**) and (**h**) air temperature (red, left axis) and relative humidity (blue, right axis), respectively, at the two sites at the time/day when stomatal conductance measurements were made. $\Phi_{PSII}$ and ETR measurements were conducted on *Cassytha pubescens*, *Acacia pycnantha* and *Bursaria spinosa* in late-February 2020 and stomatal conductance measurements were made on the two native hosts in early March 2020. Photosynthetic photon flux density (PPFD), air temperature and relative humidity were recorded with a LI-1400 data logger equipped with a quantum sensor (LI-190 SA) and relative humidity/air temperature sensor (1400-104) (LI-COR, Lincoln NEB.)

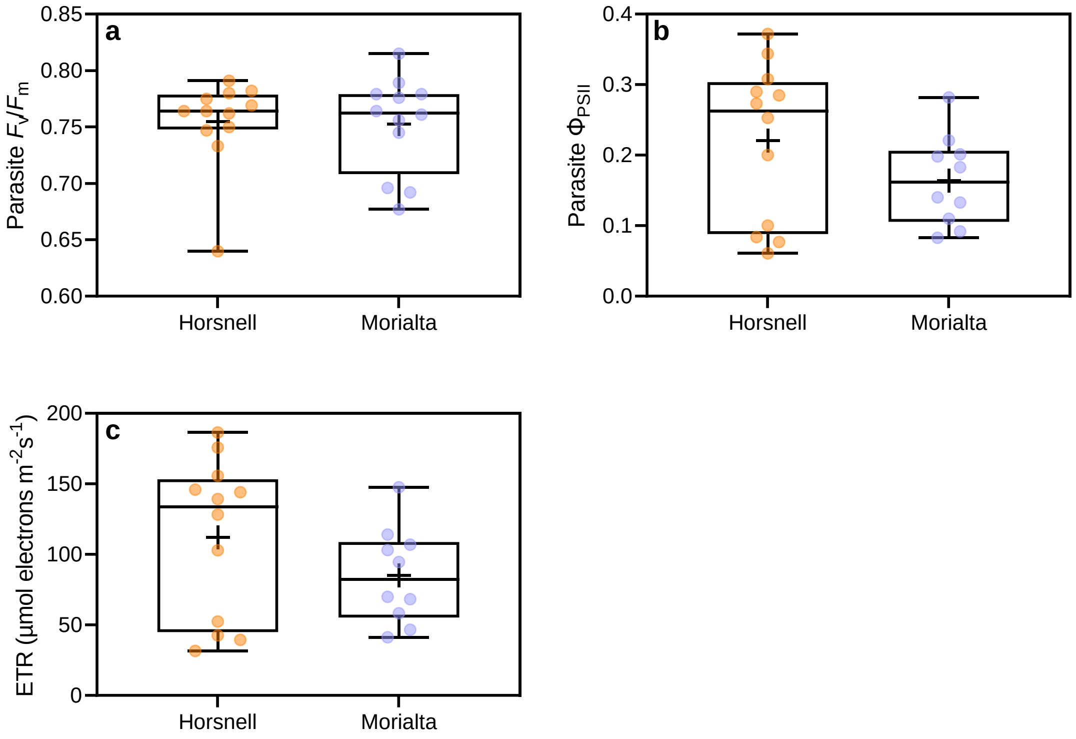


**Fig. S5** (**a**) Predawn ($F_v/F_m$) and (**b**) midday ($\Phi_{PSII}$) quantum yield and (**c**) midday electron transport rates (ETR) of *Cassytha pubescens* when infecting *Rubus anglocandicans* at two sites (Horsnell or Morialta). All data points, median, 1st and 3rd quartiles, interquartile range and mean (+ within box) are shown, different letters signify significant differences and $n$ = 12 (**a**); $n$ = 10–12 (**b**, **c**)

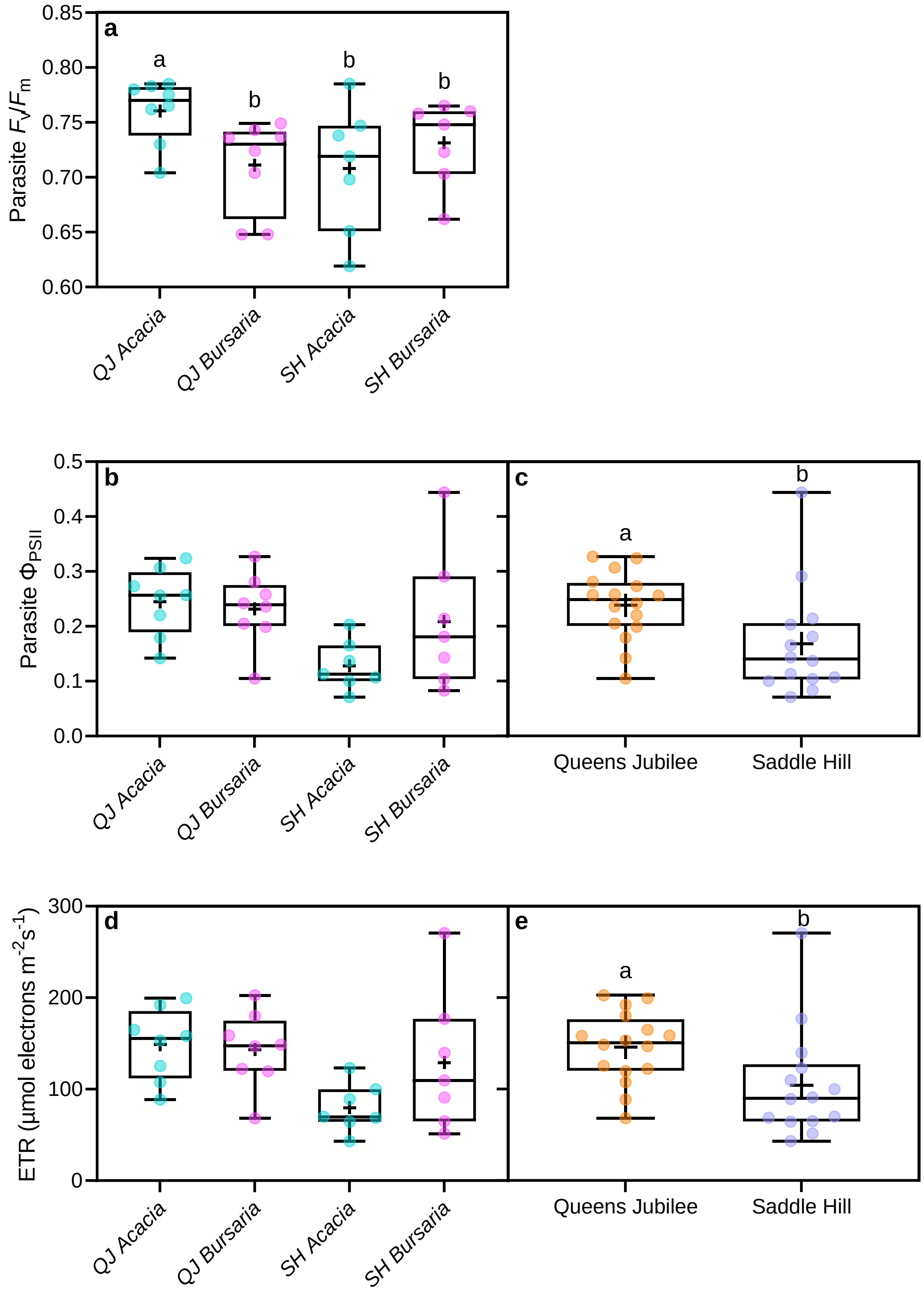


**Fig. S6** (**a**) Predawn ($F_v/F_m$) and (**b**) midday ($\Phi_{PSII}$) quantum yield and (**d**) midday electron transport rates (ETR) of *Cassytha pubescens* when infecting *Acacia pycnantha* and *Bursaria spinosa* at two field sites: Queens Jubilee (QJ) or Saddle Hill (SH). Main effect of site on parasite (**c**) $\Phi_{PSII}$ and (**e**) ETR. All sites are located in the Mt. Loft Ranges of South Australia.

All data points, median, 1st and 3rd quartiles, interquartile range and mean (+ within box) are shown, different letters signify significant differences and $n$ = 7–8 (**a**, **b**, **d**) $n$ = 14–16 (**c**, **e**)
